# Multiomics-assisted characterization of rice-Yellow Stem Borer interaction provides genomic and mechanistic insights into stem borer tolerance in rice

**DOI:** 10.1101/2023.10.10.561791

**Authors:** C. G. Gokulan, Umakanth Bangale, Vishalakshi Balija, Suneel Ballichatla, Gopi Potupureddi, Deepti Rao, Prashanth Varma, Nakul Magar, J. Karteek, M. Sravan, A. P. Padmakumari, Gouri S Laha, Subba Rao LV, Kalyani M. Barbadikar, Meenakshi Sundaram Raman, Hitendra K. Patel, M Sheshu Madhav, Ramesh V. Sonti

**Affiliations:** CSIR-Centre for Cellular and Molecular Biology, Telangana, India - 500007; ICAR-Indian Institute of Rice Research, Telangana, India - 500030; Academy of Scientific and Innovative Research, Uttar Pradesh, India - 201002; ICAR-Central Tobacco Research Institute, Andhra Pradesh, India - 533105; International Centre for Genetic Engineering and Biotechnology, New Delhi, India - 110067

**Keywords:** Rice, Yellow Stem Borer, QTL mapping, Insect resistance, Transcriptomics, Metabolomics, Markers, Plant breeding

## Abstract

Yellow stem borer (YSB), *Scirpophaga incertulas* (Walker) (Lepidoptera: Crambidae), is a major pest of rice in India, that can lead to 20-60% loss in rice production. Effective management of YSB infestation is challenged by the non-availability of adequate source of resistance and poor understanding of resistance mechanisms, thus necessitating studies for generating resources to breed YSB resistant rice and to understand rice-YSB interaction. In this study, by using bulk-segregant analysis in combination with next-generation sequencing, Quantitative Trait Loci (QTL) intervals in five rice chromosomes were mapped that could be associated with YSB tolerance at vegetative phase in a highly tolerant rice line named SM92. Further, multiple SNP markers that showed significant association with YSB tolerance in rice chromosomes 1, 5, 10, and 12 were developed. RNA-sequencing of the susceptible and tolerant lines revealed multiple genes present in the candidate QTL intervals to be differentially regulated upon YSB infestation. Comparative transcriptome analysis revealed a putative candidate gene that was predicted to encode an alpha-amylase inhibitor. Analysis of the transcriptome and metabolite profiles further revealed a possible link between phenylpropanoid metabolism and YSB tolerance. Taken together, our study provides insights on rice-YSB interaction at genomic, transcriptomic and metabolomic level, thereby facilitating the understanding of tolerance mechanism. Importantly, a promising breeding line and markers for YSB tolerance have been developed that can potentially aid in marker-assisted breeding of YSB resistance among elite rice cultivars.

**SIGNIFICANCE STATEMENT:** Global rice production is threatened by various pests, among which stem borers pose serious challenges. Hence, understanding the molecular intricacies of rice-stem borer interaction is necessary for effective pest management. Here, we used a multi-omics approach to unravel the mechanisms that might help rice combat yellow stem borer infestation, thus providing insights and scope for developing YSB tolerant rice varieties. To facilitate the latter, we developed markers that co-segregate with tolerance.

## INTRODUCTION

Several pests affect rice cultivation globally and pose serious threats to grain production and quality. Over a hundred pests have been reported to infest rice, of which 15-20 cause severe agro-economic losses (Pathak, 1968; Pathak, 1977). Yield losses caused by insects can vary depending on the insect biotype, severity of the infestation, environmental factors, growth stage and genetic background of the host (Pathak, 1977). Among the major pests of rice, stem borers pose a serious threat as they cause devastating effects on rice production in both temperate and tropical regions. Commonly found stem borers in Asia include yellow stem borer (YSB; *Scirpophaga incertulas*), striped stem borer (SSB; *Chilo suppressalis*), white stem borer, dark-headed stem borer, and pink stem borer. YSB and SSB were the cause of a steady yield loss of an estimated 5%-10% in Asia (M.D. Pathak and Z.R.Khan, 1994). YSB is a predominant pest among other stem borers in the Indian sub-continent (M.D. Pathak and Z.R. Khan, 1994; Gururaj Katti, Chitra Shanker, Padmakumari A.P., 2011). A study in a field condition revealed that YSB infestation explained about 70% variation in the yield (Gangwar *et al*., 1986). The female moths of YSB lay eggs on the leaf tips. After hatching, the larvae disperse through wind or move downwards and bore into the stem just above the water level. YSB infestation in rice is manifested as different symptoms based on the growth stage of the plants. The larvae of YSB borer into the rice stem at the base and feed on the growing tissue leading to formation of “dead hearts” in vegetative phase and white ears/ white heads in reproductive phase. Both dead hearts and white ears lead to a considerable yield loss as they both directly affect the production of rice by interfering with the number of productive tillers and panicles, respectively (Pathak, 1968; Muralidharan, K. and Pasalu, 2006). However dead heart damage at vegetative phase can be compensated up to 10% (Rubia *et al*., 1996).

The control of stem borers in the field is still challenging and is majorly dependent on timely chemical intervention. The usage of chemicals affects human health, pollutes the ecosystem, and destroys the natural predators of insects and leads to development of resistance to insecticides. Using resistant or tolerant rice cultivars to counter the attack by pests and pathogens is one of the highly recommended practices in pest management. Resistance or tolerance among different cultivars to different species of stem borers has been reported (Heinrichs, Elvis & Medrano, F.G. and Rapusas, 1985). It was observed that the resistance of these varieties was primarily due to antibiosis, that is, an antagonistic reaction of rice towards stem borer which is detrimental to the latter (Pathak, 1977). It was found that rice plants can compensate for the damage through the reallocation of metabolites and photosynthates to uninjured stems/tillers and by initiating the production of new tillers (Rubia *et al*., 1996). Compensation was observed to be an effective strategy to tolerate YSB infestation during the vegetative growth stage and not during the reproductive stage. It was proposed that breeding rice lines for tolerance to YSB based on the ability of the varieties to compensate for the YSB-caused injury could be a better approach (Rubia *et al*., 1996).

Taking note of the extent of spread and severity of the insect (Padmakumari AP and Katti G, 2018), the necessity for efficient pest management is imperative. Hence, the development of rice lines with YSB tolerance and using such lines for breeding purposes is of primary importance. This study reports the identification of a promising rice line that shows enhanced tolerance to YSB during vegetative phase of the crop. To further characterize the line and identify it as a potential source of tolerance, bulk segregant analysis and QTLseq analysis of the SM92 mapping population was performed. Putative QTL intervals linked to YSB tolerance were identified in five rice chromosomes. Using SNP genotyping of the segregating population, several markers that showed significant association with YSB tolerance were identified. In order to elucidate the mechanism of tolerance, RNA-sequencing of the tolerant and susceptible lines upon YSB infestation was performed. The analysis revealed modulation of phenyl propanoid pathway, lipid transport, and alpha-amylase inhibition could be key pathways involved in YSB tolerance in rice. Metabolite profiling further indicated accumulation of various phenylpropanoid metabolites upon YSB infestation. Overall, this work identified, characterized, and developed resources for YSB tolerance that could be of potential use in rice breeding.

## RESULTS

### SM92 was identified to be highly tolerant to YSB

A rice line named SM92 (described in Padmakumari A.P et al 2017; Potupureddi et al. 2021) was found to show tolerance to YSB during vegetative stage of growth (Figure 1A). A cut-stem assay was performed to reveal the status of YSB larvae inside the stems of SM and SM92. The results showed a significantly reduced number of live larvae inside the stems of SM92 as compared to that in SM (Figure 1B, C, D). This suggests the possibility of antibiosis mechanism in SM92.

**Figure 1.**
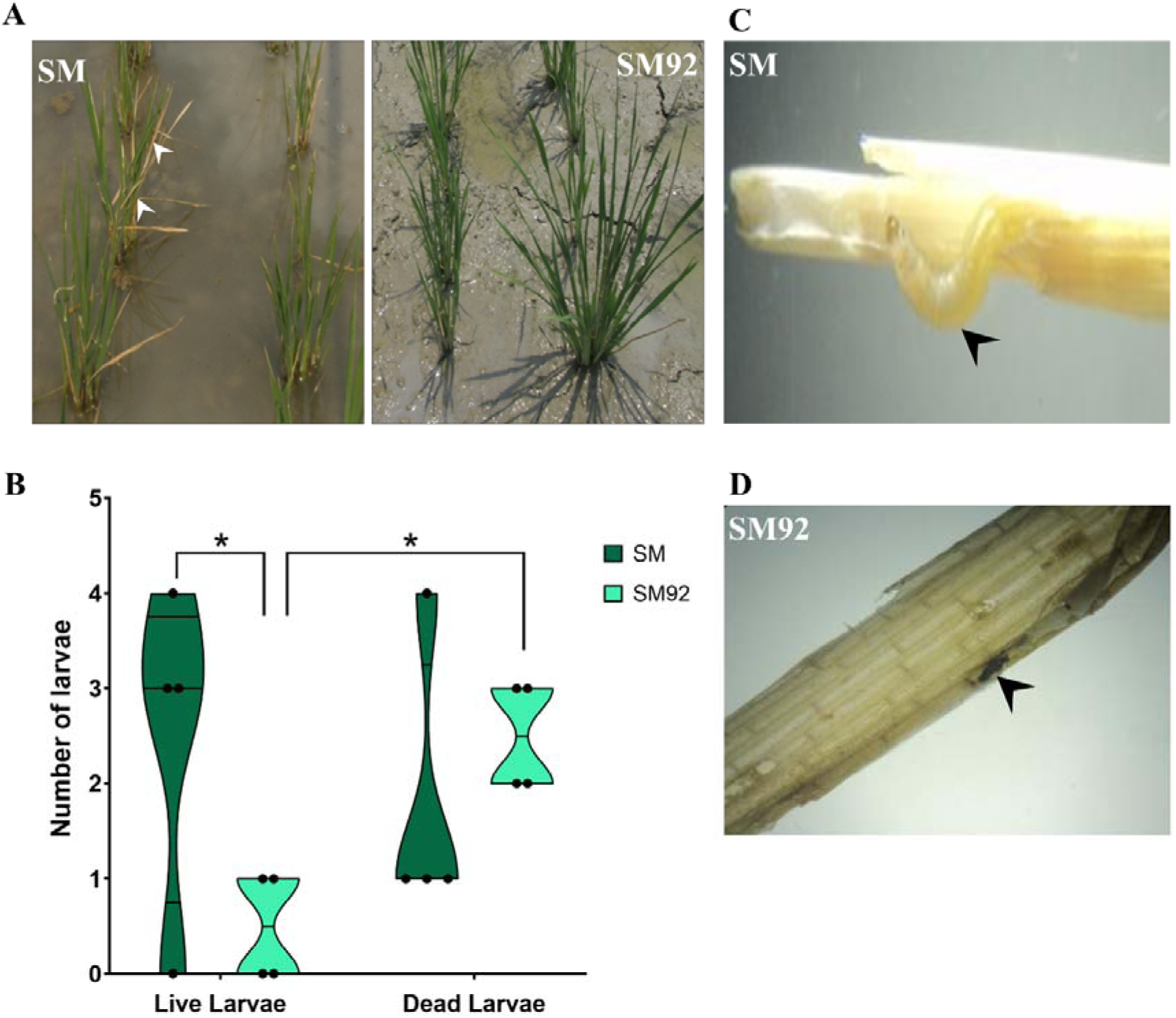
SM92 exhibited a high level of tolerance to YSB infestation. (A) Field screening with YSB larvae revealed an increased incidence of dead hearts in SM as compared to SM92. (B) Violin plot showing the number of live and dead YSB larvae in the infested stems of SM and SM92. The number of live larvae were significantly less in the stems of SM92 as compared to SM. Asterisk indicates *P*-value of less than 0.05 as calculated using a 2-way ANOVA test. (C) and (D) Images indicating the presence of a live larvae in SM and a dead larva in SM92, respectively. Black arrowheads denote the larvae.

### QTL-seq revealed putative genomic intervals linked to YSB tolerance at vegetative phase

Bulk segregant Analysis (BSA) was performed to identify the genomic regions that are associated with YSB tolerance. For this, the mutant line in the M_6_ generation (SM92) was crossed with the wildtype line. A segregating population containing 314 F_2_ progenies was screened for YSB tolerance in field conditions for dead heart symptoms (Data S1). The phenotypic segregation followed normal distribution suggesting that the trait is governed by Quantitative Trait Loci (QTL; Figure 2A). The tolerant and susceptible DNA bulks as well as the parental DNA were subjected to a high coverage (∼40X) of next-generation whole genome sequencing. The obtained raw reads were processed, aligned to the reference genome, and the variants were called. The variants were then used for the QTL-seq analysis to identify the QTL associated with YSB tolerance. QTL-seq analysis was performed using two different parameters, namely delta-SNP-index and G-prime (G’). The analyses overall revealed regions in 5 of the 12 rice chromosomes, namely chromosomes 1, 5, 7, 10, and 12, to be putatively linked to YSB tolerance (Figure 2B, C). In total, 10 QTL intervals were predicted in these 5 chromosomes (Table 1).

**Figure 2.**
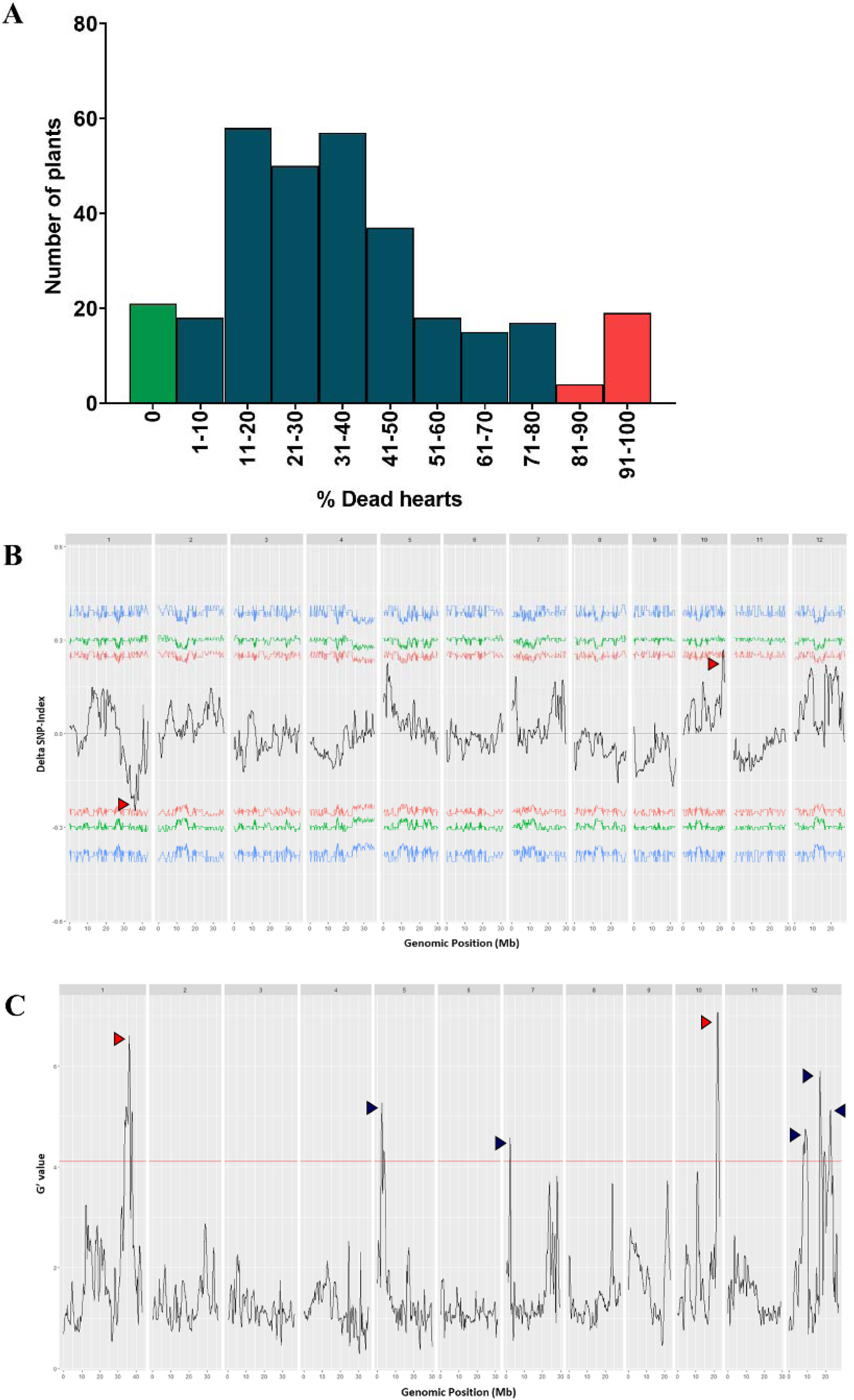
Mapping genomic loci associated with YSB tolerance in F2 population using BSA-QTL-seq. (A) Phenotype scores of the F_2_ population consisting of 314 progenies followed a continuous distribution suggesting the quantitative nature of YSB tolerance. The green bar indicates progenies that constituted tolerant bulk (n=21) and the red bars indicate the progenies that constituted susceptible bulk (n=24). (B) Delta SNP-index plot generated by the QTL-seq pipeline shows two genomic loci (red arrowheads) that could be linked to YSB tolerance. The red, green, and blue lines represent the confidence intervals at 90%, 95%, and 99%, respectively. (C) G’ plot generated by the QTL-seq pipeline shows additional genomic loci (blue arrowheads) that might be associated with YSB tolerance. The red line represents the q-value cut-off of 0.05. The black line in (B) and (C) represents the moving average line calculated using a window size of 500kb.

**Table 1:**
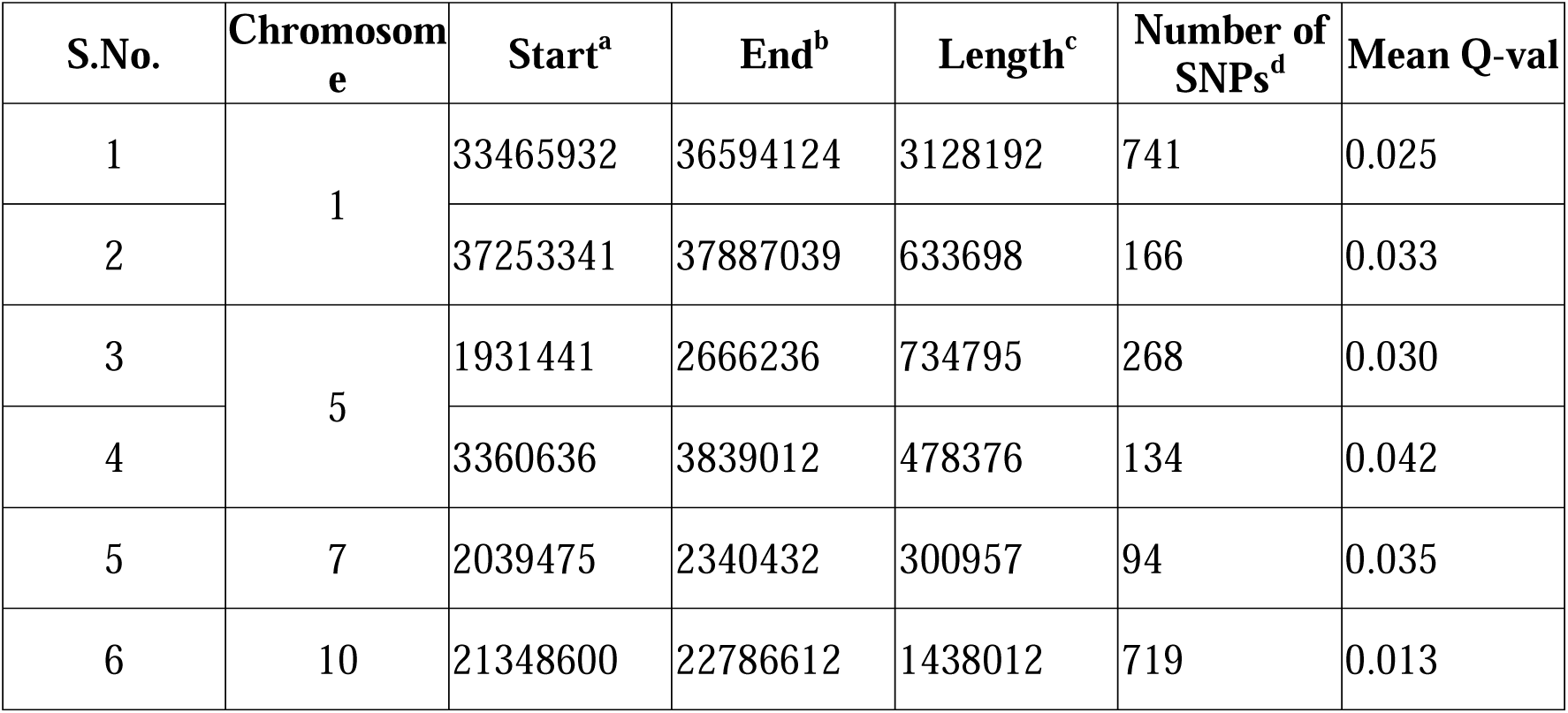
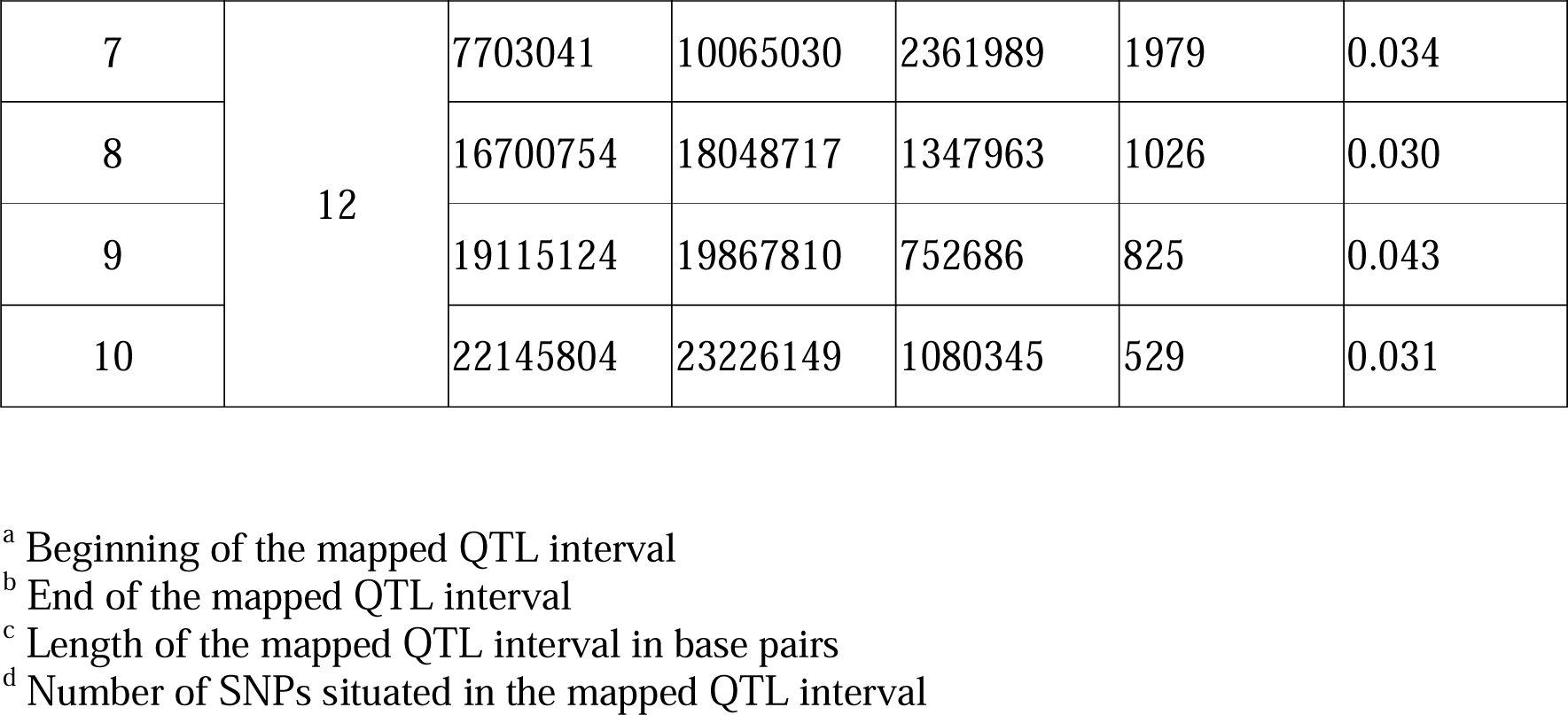
List of the putative QTL predicted to be associated with YSB tolerance. SNPs - Single Nucleotide Polymorphisms; Q-val - False Discovery Rate.

A comparison between the YSB tolerance QTL intervals and previously reported insect resistance QTL was performed. The QTL intervals identified in this study overlapped with the predicted QTL intervals linked to resistance against rice leaf folder (Chr1; Selvaraju et al., 2007) and BPH (Chr10 and Chr12; Sun et al., 2005; Jena et al., 2006) (Table S3). Although these QTL intervals are mapped for insect resistance, there is no further information on the possible genes or their alleles that might be involved in resistance. Also, no markers have been developed from these intervals for application in breeding programmes.

### A large number of genes in the QTL intervals exhibit variations

The genes that harbor SNPs and that are present in the QTL interval were annotated using Variant Effect Predictor (VEP). There were 25,065 SNPs present in the QTL intervals in all five chromosomes (Figure S1A; Data S2). About 94% of these SNPs were in non-coding regions of the genome (Figure S1B). A total of 1101 genes harbor SNPs which include SNPs in coding (UTRs and exonic) and/or non-coding (upstream, downstream, and intronic) regions. Genes harboring SNPs in their coding regions were 459 in number and 34 among them harbored SNPs that were predicted to be deleterious (Data S3). Over-representation analysis of the genes that harbor mutations in the coding sequences revealed a significant enrichment of laccase-encoding genes, that are involved in phenylpropanoid/lignin metabolism (Figure S1C).

### Marker-Trait Association Analysis revealed markers exhibiting significant association with YSB tolerance at vegetative phase

A set of fifty SNPs were selected as markers for genotyping the F_2_ individuals using Kompetitive Allele Specific PCR (KASP^TM^, LGC Genomics, UK; Data S4, S5). These SNPs were selected based on their effect on the genes that harbor them, the genomic distance between the preceding and the following SNP in such a way that the distance between the two markers is less than 250-300 kbp. KASP assay was performed from the F_2_ plants showing extreme as well as intermediate phenotypes (Data S6). Further, a marker-trait association analysis was carried out to identify the markers that are significantly associated with the phenotype (Data S7). The KASP-based genotyping revealed a significant (*P*<0.05) association of fourteen alleles from chromosomes 1, 5, 10, and 12 with YSB tolerance in the F_2_ individuals (Table 2). Each of the regions was calculated to explain about 5% to 9% phenotype variation observed in the F_2_ progenies.

**Table 2:** Markers that are significantly associated with YSB tolerance.

| Marker | Chromosome | Position | REF<sup>a</sup> | ALT<sup>b</sup> | SNP-index Tol<sup>c</sup> | SNP-index Sus<sup>d</sup> | Delta SNP-index | P | % PVE | Gene ID | Description |
| --- | --- | --- | --- | --- | --- | --- | --- | --- | --- | --- | --- |
| S2 | 1 | 33886610 | C | G | 0 | 0.61 | -0.61 | 0.034 | 5.7 | Os01g0800266 | Similar to Phosphoglycerate kinase. (Os01t0800266-00) |
| S3 |  | 33957364 | G | A | 0.29 | 0.62 | -0.33 | 0.022 | 6.7 | Os01g0801700 | Ribosomal protein S35, mitochondrial domain containing protein. (Os01t0801700-01) |
| S5 |  | 34710185 | G | A | 0 | 0.63 | -0.63 | 0.007 | 9.2 | Os01g0816400 | Similar to microtubule organization protein. (Os01t0816400-01) |
| S6 |  | 35088627 | C | T | 0 | 0.69 | -0.69 | 0.019 | 7.1 | Non-exonic<sup>†</sup> | - |
| S17 | 5 | 2366109 | G | A | 0.65 | 0 | 0.65 | 0.047 | 5.0 | Os05g0141200 | Hypothetical conserved gene. (Os05t0141200-00) |
| S18 |  | 3157367 | C | T | 0.59 | 0 | 0.59 | 0.033 | 5.8 | Non-exonic | - |
| S19 |  | 3743040 | G | T | 0.91 | 0 | 0.91 | 0.022 | 6.6 | Non-exonic | - |
| S20 |  | 4321188 | C | T | 0.66 | 0 | 0.66 | 0.047 | 5.1 | Os05g0172300 | Domain of unknown function DUF632 domain containing protein. (Os05t0172300-01) |
| S28 | 10 | 21786052 | C | T | 0.73 | 0 | 0.73 | 0.017 | 7.2 | Non-exonic | - |
| S29 |  | 21853916 | T | A | 0.82 | 0 | 0.82 | 0.017 | 7.2 | Non-exonic | - |
| S30 |  | 21955122 | C | T | 0.68 | 0 | 0.68 | 0.017 | 7.2 | Os10g0557800 | Conserved hypothetical protein. (Os10t0557800-01) |
| S31 |  | 21955776 | T | A | 0.76 | 0.45 | 0.31 | 0.038 | 5.5 | Os10g0557900 | Peptidase M10A and M12B, matrixin and adamalysin family protein. (Os10t0557900-00) |
| S32 |  | 21965877 | C | G | 0.69 | 0.38 | 0.31 | 0.017 | 7.2 | Os10g0558125 | Similar to Matrixin family protein, expressed. (Os10t0558125-00) |
| S45 | 12 | 18013596 | A | T | 0.63 | 0 | 0.63 | 0.034 | 5.8 | Non-exonic | - |

<sup>a</sup> Allele present in the wildtype line
<sup>b</sup> Allele present in the mutant line
<sup>c</sup> SNP-index of the tolerant bulk
<sup>d</sup> SNP-index of the susceptible bulk
\* Control – SNPs from genomic regions outside QTL intervals
<sup>†</sup> Non-exonic – SNPs from the QTL intervals that are not exonic
%PVE - % Phenotypic Variation Explained

### YSB infestation causes an extensive transcriptional reprogramming

Transcriptomics-based characterization of rice-YSB interaction was performed to understand the possible mechanism of tolerance to YSB. For this, RNA sequencing (RNA-seq) was performed on RNA isolated from the stem tissues of SM and SM92, with and without YSB infestation (3 days post infestation). The analysis of RNA-seq data showed several differentially expressed genes (DEGs) with overlapping and unique DEGs between SM and SM92 (Figure 3A, B; Data S8, S9). The number of DEGs was slightly more in SM (n=1950) when compared to SM92 (n=1857). Also, SM92 had a lesser number of downregulated genes than SM. Overall, only 13.7% of the DEGs were common between SM and SM92, suggesting a contrasting transcriptional reprogramming occurring in tolerant and susceptible lines upon YSB infestation.

**Figure 3.**
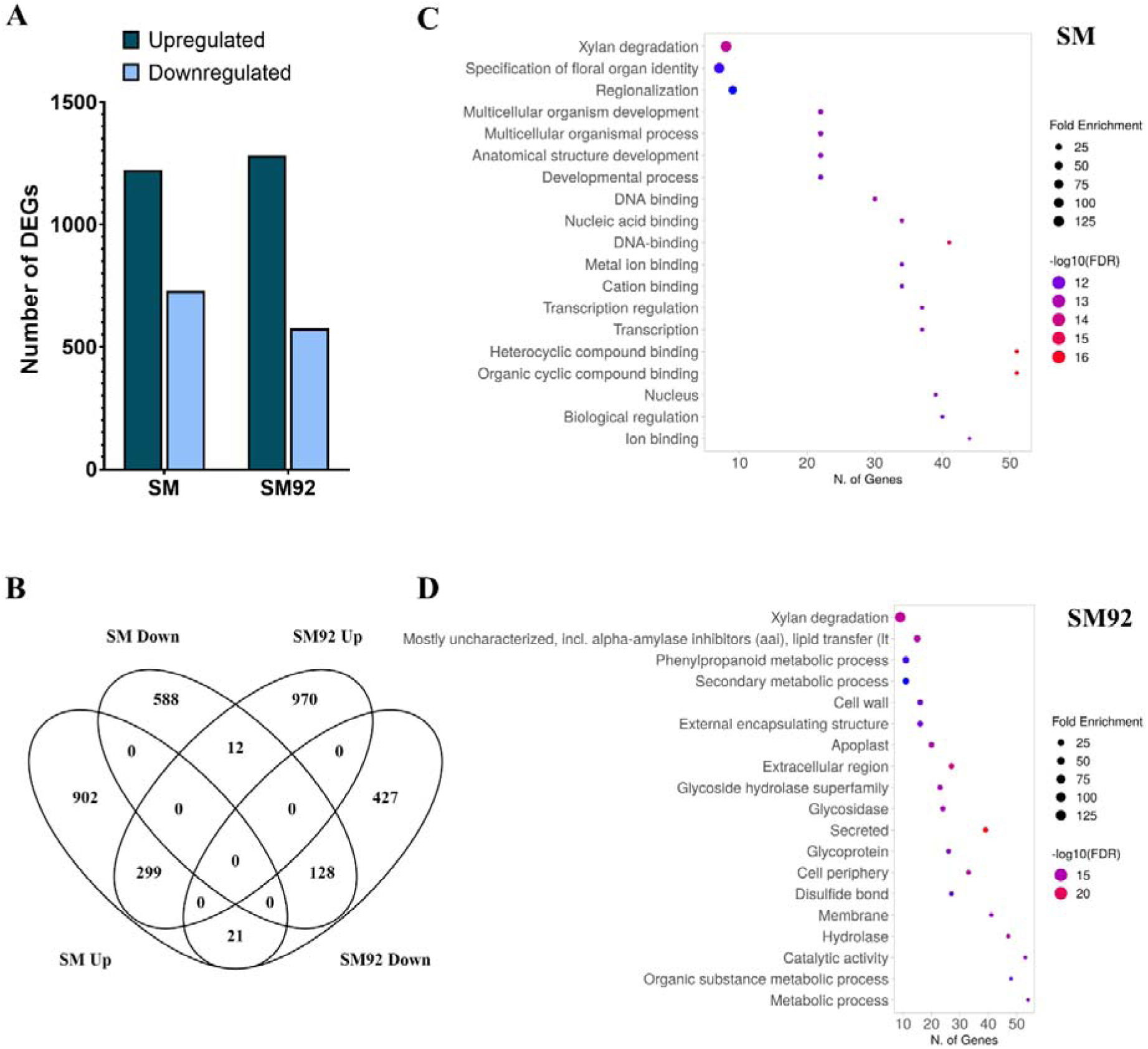
Transcriptome profiling revealed putative rice functions associated with YSB tolerance. (A) Bar chart depicting the number of differentially expressed genes in SM and SM92 upon YSB infestation in comparison to uninfested samples, as obtained through RNA-sequencing based transcriptome profiling of stem samples at 72 hours post infestation. (B) Venn diagram comparing the differentially expressed genes in SM and SM92. (C) and (D) Gene enrichment analysis showing the upregulated pathways in SM and SM92 post YSB infestation.

Gene ontology (GO) and pathway enrichment analysis of the DEGs revealed the similarities and differences between SM and SM92 in terms of their transcriptomics response to YSB infestation (Figure 3C, D). Specifically, YSB-induced genes in SM92 were enriched in phenylpropanoid metabolism, alpha-amylase inhibitors/lipid transfer protein and glycoside hydrolases (Figure 3D). An in-depth analysis using MapMan revealed genes belonging to the GO term ribosome biogenesis were exclusively downregulated in SM (Figure S2A). The categories consisting of genes involved in the metabolism of simple phenols, lignin and lignans showed a striking difference between SM and SM92, wherein these genes were majorly upregulated in SM92 (Figure S2B).

### Phenylpropanoid pathway genes show remarkably contrasting expression pattern at vegetative phase infestation in tolerant and susceptible lines

Genes involved in the biosynthesis of phenylpropanoids showed a remarkable upregulation in SM92 while some of the genes were downregulated in SM (Figure 3D; Figure S3). The Phenylpropanoid Pathway (PPP) is an important secondary metabolism pathway that is essential for normal plant growth, development and defense against various pests and pathogens (Dong and Lin, 2021). PPP leads to the production of various types of lignin and lignin derivatives. In line with this, overall, thirteen *laccase*-encoding genes, that are predicted to be involved in lignin metabolism, were upregulated in SM92 upon YSB infestation. Markedly, five of the thirteen *laccase* genes are present within the chromosome 1 QTL interval (Table S4). Further analysis revealed that cumulatively 118 genes that are in the QTL intervals are differentially expressed in SM and SM92 upon YSB infestation (Data S10). A deeper look into these genes indicated lignin metabolism-related genes, *laccases* (n=5), to be enriched among the genes upregulated exclusively in SM92 (Figure S4A). Promoter analysis of the five *laccase* genes revealed the presence of tandem CATG palindromic motif (CATGCATG) in the promoters of all five genes (Fig S4B). Further analysis indicated a significant overlap between the observed motif and the binding element of a B3-domain containing transcription factor (Figure S4C). It could hence be speculated that a common mode of regulation might be acting on these *laccase* genes in SM92, probably mediated by a B3-domain containing transcription factor.

### Comparative analysis suggests putative candidate genes for stem borer tolerance in rice

A comparative transcriptomics analysis was performed between the data generated in this study and a previously published data where rice lines with varying degree of resistance to striped stem borer (SSB) were infested with SSB (Wang *et al*., 2018). It was observed that 527 genes were commonly upregulated by SSB and YSB in the respective tolerant lines (Figure 4A; Data S11). A gene set enrichment analysis revealed that genes annotated as alpha-amylase inhibitors/lipid transfer protein, carbohydrate metabolism, and cell wall-related genes were significantly enriched (Figure 4B). A striking observation was that the highest upregulated gene in SM92 upon YSB infestation – *OsLTPL146*, a Lipid Transfer Protein-like (LTPL) gene – is present within the mapped QTL interval in Chromosome 10. In addition, the same gene was significantly downregulated in SM. *OsLTPL146* exhibited a similar expression pattern upon SSB infestation as well. That is, *OsLTPL146* was upregulated in a resistant cultivar and downregulated in a susceptible cultivar upon SSB infestation (Figure 4C). Interestingly, it was also observed that twenty-five lipid-metabolism/transport related genes majorly including *LTPL* class of genes were upregulated in SM92 upon YSB infestation, while only five genes showed differential expression in SM (Figure S5A). Other lipid/fatty-acid metabolism related genes that are involved in cutin, suberine, and wax synthesis were also exclusively upregulated in SM92 (Figure S6). Another unique expression pattern observed was that of the glutathione metabolism genes consisting of the Phi and Tau classes of *glutathione transferases* (*GSTs*). These genes were exclusively upregulated in SM92 post YSB attack (Figure S6).

**Figure 4.**
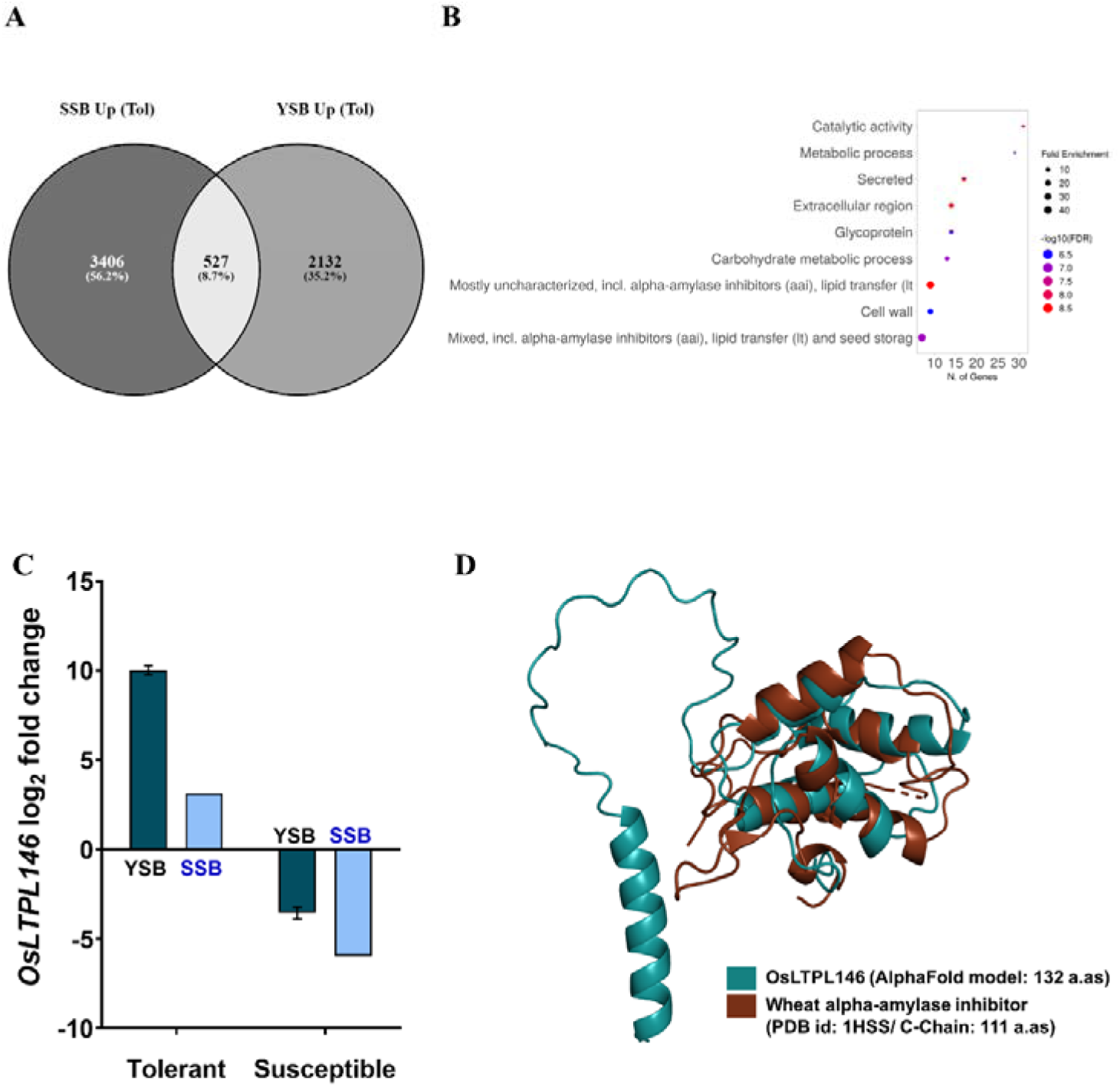
Comparative transcriptome indicated a possible candidate gene for YSB tolerance. (A) Venn diagram showing the comparison of genes upregulated in rice lines that are tolerant to striped stem borer (SSB) from Wang et al (2018) and yellow stem borer (YSB). The intersection indicates 527 genes that are commonly upregulated. (B) Gene enrichment analysis of the stem borers-induced genes revealed the enrichment of functions involved in alpha-amylase inhibition, glycoproteins, and secreted proteins. (C) Expression of the gene *OsLTPL16* was observed to be upregulated by SSB and YSB in tolerant rice lines and downregulated in susceptible rice lines. (D) Structure prediction and homology search suggests that *OsLTPL146* might encode an alpha-amylase inhibitor. Structure model of OsLTPL146 is colored green and the solved structure of wheat alpha-amylase inhibitor (PDB:1hss) is colored brown.

As mentioned above, twenty-five enriched genes belonged to the lipid metabolism/transport category. Of the twenty-five genes, twenty-two genes were annotated as “Protease inhibitor/seed storage/LTP family protein precursor”. To elucidate the actual biochemical properties of these gene products, the structures of all the twenty-five protein sequences were predicted using AlphaFold2 and their structural homologues were identified. The results suggest that fifteen genes likely encode lipid transfer proteins as their structures were homologous to characterized lipid transfer proteins from various organisms, five gene products were homologous to alpha-amylase inhibitors and of the remaining five genes, one coded for a lyase, one for a phospholipase D, one for annexin and the structures could not be predicted for the other two gene products (Figure S5B).

Notably, structure prediction analysis suggests that the predicted structure of *OsLTPL146* is homologous to an alpha-amylase inhibitor whose structure is experimentally solved. This result indicates that *OsLTPL146* – the highest upregulated gene in SM92 upon YSB infestation – likely encodes an alpha-amylase inhibitor (Figure 4D). Further, owing to the difference in the expression pattern of *OsLTPL146* in SM and SM92, the promoter sequence of *OsLTPL146* in SM and SM92 were compared. The alignment revealed considerable variations in the promoter and 5’ untranslated region (UTR) of the gene between SM and SM92in certain known cis-elements. The variations included single nucleotide polymorphisms, small insertions, and deletions (Figure S7). Whether the variations in the promoter/regulatory sequence of *OsLTPL146* are involved in the YSB-mediated induction of the transcript and/or YSB tolerance is another avenue for further research. Taken together, detailed investigation of OsLTPL146 promoter variation and protein biochemistry could reveal its role in stem borer resistance.

### Levels of a few phenylpropanoid pathway metabolites are altered upon YSB infestation

An untargeted metabolite profiling of sheath samples that were either uninfested or infested with YSB larvae was performed to gain insights into the mechanism of tolerance to YSB. Over 1000 metabolites were identified in all the tested conditions (Data S12, and S13). The overall profile indicated a difference in the metabolite levels between uninfested SM and SM92 plants. However, YSB infestation seems to result in highly comparable metabolite profile in both SM and SM92 (Figure 5A). Further, the levels of phenylpropanoid pathway (PPP) associated metabolites were analysed specifically, as the RNA-seq data indicated an upregulation of genes associated with the pathway. The results showed the presence of 27, and 25 PPP metabolites in SM and SM92 samples, respectively (Figure S8). Four of the PPP metabolites including 5-O-Caffeoylshikimic acid, Coniferyl acetate, Caffeate, and L-Phenylalanine exhibited significant differential accumulation in SM and/or SM92 upon YSB infestation when compared to uninfested samples (Figure 5B). YSB infestation resulted in increased levels of 5-O-Caffeoylshikimic acid and Coniferyl acetate in both SM and SM92. On the other hand, Caffeate and L-Phenylalanine exhibited significantly reduced accumulation in SM92 upon YSB infestation. Owing to the variation among the samples, difference in the levels of other metabolites could not be reliably identified. Targeted analysis of PPP metabolites could provide a better understanding of the regulation of the pathway with respect to YSB infestation.

**Figure 5.**
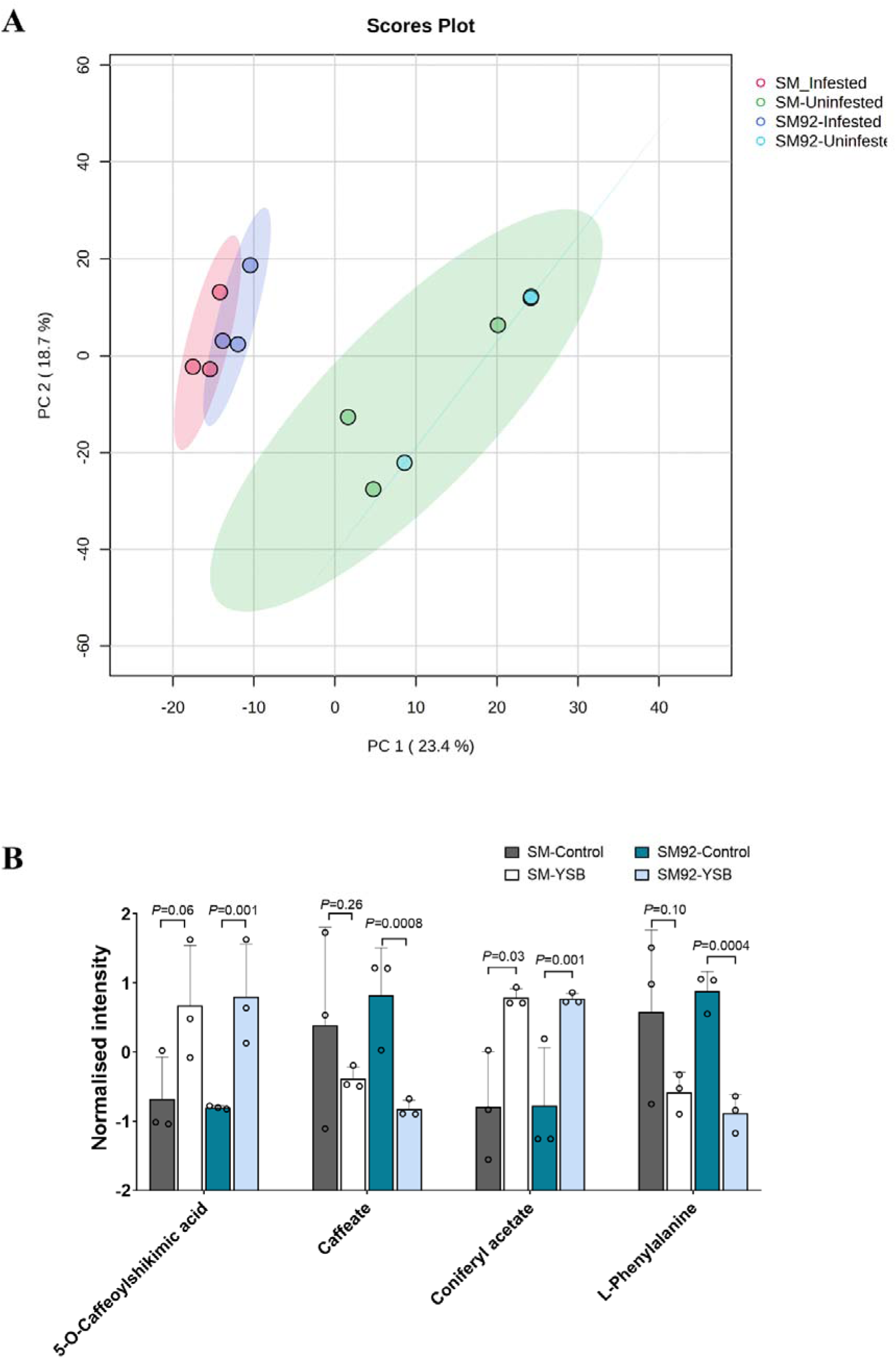
YSB infestation causes considerable changes in the plant metabolite composition. (A) Principal Component Analysis (PCA) plot showing the clustering of SM and SM92 samples infested with YSB (pink- and purple-shaded oval) indicating an overlapping response in both SM and SM92. The uninfested samples, however, showed a loose clustering, probably indicating differences in the basal metabolite composition. (B) Box plot showing the normalized intensity of certain phenylpropanoid pathway metabolites in SM and SM92 upon YSB infestation and uninfested conditions. *P-*values were calculated using unpaired t-test with multiple comparison correction using Holm-Sidak method. P<0.05 is considered to be statistically significant.

## DISCUSSION

Stem borers cause maximum yield loss among all the insects that attack rice in India (Savary *et al*., 2019). Complete resistance to stem borers in nature is scarce and unidentified yet. Being a monophagous pest, it feeds only on rice plants and perpetuates. As a borer, by nature of its feeding habit it must feed on the plant for its survival. The damage occurs in the process of entering the plant and therefore some extent of damage is inevitable. Hence, the dependance on chemical insecticides prevail despite their negative impact on health and the ecosystem (Su *et al*., 2014). Most stem borer tolerant varieties of rice exhibit compensation and antibiosis traits (Rubia *et al*., 1996). However, the exact mechanism of tolerance and the molecular players involved are unknown.

The advent of bulk segregant analysis and next-generation sequencing methods have accelerated QTL identification and gene mapping for the traits of interest. This study identified a rice line, named SM92, that is highly tolerant to YSB. Screening the F_2_ population, generated through bi-parental mating, at vegetative phase revealed that YSB tolerance to be a quantitative trait. Therefore, the segregants with extreme phenotypes were used for preparing tolerant and susceptible DNA bulks and were subjected to high-coverage sequencing. Analysis of the sequence data using QTL-seq pipeline indicated the presence of QTL intervals in five rice chromosomes that are putatively linked to YSB tolerance, thus validating the observed segregation pattern. Notably, three of the identified genomic regions were found to be overlapping with known insect resistance QTL intervals (Table S3). That is the Chr1 QTL was partly overlapping with a previously identified rice leaf folder resistance QTL whereas the Chr10 and Chr12 QTL intervals partly overlap with BPH resistance QTL. This observation suggests that the identified genomic intervals are of potential importance in insect resistance/tolerance and are likely to contain genes/alleles that are part of the insect resistance/tolerance mechanism of rice. The QTL mapping data indicated that SM alleles contribute to tolerance arising from Chromosome 1 QTL interval whereas SM92 alleles contribute to tolerance arising from other intervals (Figure 2B). Further analysis indicated that *laccase* genes, involved in lignin metabolism, are overrepresented in the QTL intervals.

### Laccases and plant defense

Laccase (LAC; EC1.10.3.2) belongs to a multi-copper oxidase family of enzymes that play a crucial role in plant development and stress responses (Yu *et al*., 2021). Various studies have identified the multi-faceted roles of laccases in plants that include cell wall elongation, secondary cell wall formation, lignification, pigmentation, flavonoid biosynthesis, metal ion stress, and seed setting. Laccases are involved in catalyzing the polymerization of lignin monomers in the apoplast (Zhao *et al*., 2013). Lignin, being a component of the secondary cell wall, plays important roles in protecting cells from pathogens. Previous studies have established a positive correlation between lignin accumulation and resistance to bacterial and fungal infections (Lee *et al*., 2019; Peltier *et al*., 2009). Previous studies have also associated the levels of lignin pathway precursors including hydroxycinnamic acid positively with resistance to stem borers (Santiago *et al*., 2013; Malvar *et al*., 2008). A study in maize showed a positive correlation between levels of phenylpropanoids including *p*-coumarate and syringyl lignin to Mediterranean Corn Borer resistance (Gesteiro *et al*., 2021). Our transcriptome profiling indicated thirteen *laccase* genes to be significantly upregulated in SM92 upon YSB infestation. Additionally, some of the genes involved in the phenylpropanoid metabolism pathway exhibited a strikingly contrast expression pattern in SM (downregulated) and SM92 (upregulated) upon YSB infestation. Notably, five of the thirteen *laccase* genes are in the identified QTL intervals in Chromosomes 1. It was also noted that some of the phenylpropanoid pathway genes were downregulated in SM, the susceptible line (Figure S3). Together, the data suggest a likely involvement of lignin metabolism in conferring tolerance to YSB. Although previous studies have pointed out a generic role of laccases in pathogen/pest resistance, our study has provided promising candidate genes that can be studied further to understand the mechanism of tolerance at molecular level.

### Ribosomal proteins as potential candidates to understand plant defense mechanisms

On examining the gene enrichment in the susceptible interaction, a remarkable number of genes that code for ribosomal subunit proteins (RPs – Ribosomal proteins) were found to be significantly downregulated. Both the large and small subunit protein-encoding genes were highly downregulated. Previous studies have pointed out that RPs possess moonlighting functions. Such extra-ribosomal functions of RPs include RNA-binding and promoting/preventing translation, role in apoptosis, cell-cycle regulation, and stress resilience in prokaryotes as well as eukaryotes (Wool, 1996; Weisberg, 2008; Warner and McIntosh, 2009; Trautmann and Ramsey, 2022). In plants, two RPs viz. RPL12 (Ribosomal Protein Large subunit 12) and RPL19 in *Nicotiana benthamiana* and *Arabidopsis thaliana* were shown to be important for non-host resistance as the mutants exhibited delayed hypersensitive response to non-host pathogens (Nagaraj *et al*., 2016; Ramu *et al*., 2020). A targeted transcriptome profiling of thirty-four large subunit genes indicated that they are highly stress responsive. The genes were either upregulated or downregulated depending on the nature of the stress. Infection of rice plants with the bacterial blight pathogen caused upregulation of the majority of the tested RPs (Moin *et al*., 2016). Downy mildew infection in grapevine resulted in accumulation of RPs during the early timepoints post-infection (Santos *et al*., 2020). However, our data revealed a significant downregulation of ribosomal protein-coding genes. This could be an energy-conserving strategy that the host deploys to enhance defense against insect attack. But it is also possible that reduced expression of RPs is a consequence of the mechanisms that the pest uses to suppress defense, and this ultimately results in the host succumbing to the attack. It is noteworthy that the tolerant line did not show any alteration in the expression of RP-encoding genes (Figure S2A), thus suggesting susceptibility being a consequence of RP gene downregulation. Further studies that follow the expression pattern of RP-encoding genes at different time points post infestation can provide a clearer picture of the role of RPs in rice-stem borer interaction.

### Insights gained through structure prediction analysis

Another striking observation from the transcriptomics data was the differential expression of approximately twenty genes that were annotated as Lipid transfer-like proteins (LTPLs)/Protease inhibitors (PIs). One of the genes *OsLTPL146* showed the highest upregulation (log_2_FC>10) upon YSB infestation in the tolerant line while it was downregulated in the susceptible line. Importantly, *OsLTPL146* is located in the Chromosome 10 QTL interval, suggesting a possible link between the expression of the gene and YSB tolerance at vegetative phase. A similar pattern of expression was observed for *OsLTPL146* in an independent transcriptomics study performed in rice upon striped stem borer infestation (Wang *et al*., 2018). Owing to the high sequence similarity between LTPs and PIs, based on the annotation and sequence alone it is hard to predict the biochemical function of their gene products. Hence, the recently developed, artificial intelligence-based AlphaFold2 was used to predict the structure of the differentially expressed LTPLs (Jumper *et al*., 2021). The results indicated that fifteen out of the twenty LTPLs exhibit structural homology to lipid transfer proteins while the rest five were homologous to alpha-amylase inhibitors. The presence of proteins of different classes suggest that multiple modes of tolerance are likely in action including protease inhibition, lipid transfer, and alpha-amylase inhibition.

Lipid transfer proteins (LTP and LTP-like (LTPL) proteins) have been associated with pathogen resistance, plant development, abiotic stress development, and cuticular wax metabolism among others (Gao *et al*., 2022). LTPs are small proteins that bind to lipids and aid in their transport across cellular compartments (Salminen *et al*., 2016). Multiple rice LTPs/LTPLs have been associated with pollen development, tolerance to abiotic stress such as drought, and plant development (Gao *et al*., 2022). *OsLTP5* was found to be induced by cutin monomers, abscisic acid, and salicylic acid (Kim *et al*., 2008). Cutin monomers possess a defense elicitation property against a potential invasion (Schweizer *et al*., 1996; Fauth *et al*., 1998; Kolattukudy *et al*., 1995). Class I LTPs were identified to be induced upon infection by the blast fungus, *Magnaporthe oryzae* (Tae *et al*., 2006). The damage caused by fungus and chewing insects results in cuticle damage which in turn leads to the generation of cutin monomers that subsequently activate defense signaling such as accumulation of H_2_O_2_ (Fauth *et al*., 1998). Moreover, our transcriptome analysis also revealed enrichment of genes involved in cutin/wax biosynthesis to be upregulated in SM92 upon YSB infestation, suggesting a possible production of cutin and/or reorganization of the cuticle being associated with YSB tolerance.

Alpha-amylase inhibitors (AAIs) belong to multi-family proteins that interact with and inhibit alpha-amylase enzymes (Franco *et al*., 2002; Oneda *et al*., 2004). Alpha amylases are important digestive enzymes that help in the digestion of plant-derived starch/amylose that are ingested by the insects (Lage, 2018). The evolutionary arms race between plants and pests has resulted in the evolution of AAIs that inhibit alpha-amylase activity in the insect gut and thus negatively impacts insect survival (Franco *et al*., 2002).

### Multi-layered mechanisms to counter stem borer infestation in rice

Studies so far have identified such tolerance-associated mechanisms individually in different plants/varieties. But our observations in combination with other reports in the literature suggest a multimodal mechanism of tolerance to YSB in SM92. Our results suggest that the striking upregulation of multiple Lipid Transfer Protein like genes and alpha-amylase inhibitor genes in SM92 is potentially linked to YSB tolerance in a manner that is either dependent on or independent of wax/cuticle damage. Upregulation of a considerable number of lipid transfer protein-like (LTPL) genes indicates a possibility of major alteration in the lipid composition of the cells. The upregulation of cutin/wax synthesis genes in combination with *LTPLs* provides a reasonable indication of the involvement of lipid translocation during insect attack as a plant defense strategy. More studies on this can provide novel insights into plant defense signaling and mechanisms involving lipid moieties. Also, future studies directed towards OsLTPL146, and its amylase inhibition activity could provide a native source of alpha-amylase inhibition thereby eliminating the need for a source of foreign origin.

Additionally, the upregulation of phenylpropanoid metabolism genes and laccases adds one more layer to the complex mechanism of YSB tolerance in SM92. The presence of signature motif in the promoter sequence of similarly regulated *laccases* suggests a common regulatory network that might control the expression of lignin metabolism genes in SM92 upon YSB infestation. Therefore, SM92 is a very promising line for gaining insights into fundamental resistance mechanisms as well as for breeding programmes to introgress YSB tolerance into breeding lines. The former can be accomplished by studying the candidate genes from this study based on QTL-seq and transcriptomics results, while the latter has been facilitated by the markers that were identified to be significantly associated with tolerance.

### Breeding for YSB tolerance

Marker-assisted breeding is a viable option for introgressing traits from one crop variety to another. In this regard, several SNP markers have been identified in this study that span the QTL intervals in four different chromosomes. Notably, the Chr10 QTL interval was fine mapped to a ∼180kb region with five linked SNP markers. On the other hand, a few markers have been identified to be linked to YSB tolerance in Chr1, Chr5, and Chr12 QTL intervals. Further mapping studies and marker-trait association analyses using high density markers could help in fine mapping the QTL intervals thereby providing additional markers with tight linkage/association to YSB tolerance.

Overall, to the best of our knowledge, the study presented here is the first to provide important insights at genomic, transcriptomic, and metabolomic levels to uncover the genomic loci and mechanisms of tolerance to yellow stem borer in rice. Stem borers, one of the major group of pests affecting global rice production, in combination with changing and erratic climatic conditions, pose a serious threat to food security. Knowledge from this study would aid in orienting future research to better understand the mechanisms as well as provide additional gene targets to breed stem borer resistance in rice and possibly in other related crops.

## EXPERIMENTAL PROCEDURES

### Plant materials and Insect

Plant materials used for this study include the cultivar Samba Mahsuri (SM) and the EMS mutant line named SM92. SM92 is registered as a germplasm at ICAR-National Bureau of Plant Genetic Resources (National Identity: IC646828). Yellow Stem Borer (YSB; *S. incertulas*) moths were collected from the rice field and were released for egg laying on rice plants. The neonate larvae emerging from the egg masses were used for infestation of the mutant lines, F_2_, and F_2:3_ progeny under field conditions by releasing 2 neonate larvae per tiller at the maximum tillering phase (Bentur JS *et al*, 2011; Padmakumari AP and Katti G, 2018). The phenotype was scored when the dead heart symptoms were apparent on the susceptible cultivar (IRRI, 2014). All the YSB screening activities were conducted in the greenhouse and/or rice field facility of ICAR – Indian Institute of Rice Research, Hyderabad, India (17.32LJN, 78.39LJE).

### Cut-stem assay

Cut stem insect bioassays were carried out at the ICAR-IIRR. At vegetative phase, stem cuttings were collected from healthy tillers of both SM and SM92. The stems were cut into 4-5 cm long pieces and placed on Petri plates containing Whatman paper moistened with benzimidazole (75 mg/L). Freshly hatched neonate YSB larvae were then released onto each cut stem at 5 larvae/stem. The plates were kept in an incubator at 27±2°C. After 4 days, the plates were opened, and the stems were dissected carefully under a microscope. Before dissection, the stem pieces were observed for the presence of feeding holes. Upon dissection, the stems were observed for feeding injury and for the number of dead/surviving larvae were counted.

### Bulk segregant Analysis and sequencing

For mapping the genomic region(s) associated with YSB tolerance, bulk segregant analysis (BSA) followed by next-generation sequencing and QTL-seq analysis was performed as described earlier (Takagi *et al*., 2013; Mansfeld and Grumet, 2018). A cross was made between SM (WT; recipient) and SM92 (Mutant; donor). The F_1_ plants were confirmed for hybridity and advanced to F_2_ generation comprising 314 plants. The F_2_ progenies and the parental lines were screened during the vegetative growth phase. The plants were scored based on the percentage of dead heart as recommended by the Standard Evaluation System of Rice (IRRI, 2014). F_2_ plants showing the extreme phenotypes i.e., tolerance (n=21) and susceptibility (n=23) to YSB were selected to constitute the tolerant and susceptible bulks, respectively, after confirming the phenotypes of the F_2:3_ lines. DNA from the bulk constituents and parents were isolated using the CTAB method and quantified using Qubit^TM^ High-Sensitivity dsDNA Assay in a Qubit^TM^ 4 fluorometer (Thermo Fisher Scientific, USA). The DNA quality was assessed on 0.8% agarose gel through electrophoresis. Equal concentrations of DNA from the tolerant and susceptible plants were bulked separately. 1 µg DNA from all the samples was used to prepare sequencing libraries with TruSeq Nano DNA HT Sample Preparation Kit (Illumina, USA) and qualified libraries were sequenced on the Illumina HiSeq2500 platform to obtain 2x150 bp reads at approximately 40x coverage. The library preparation and sequencing were outsourced to Nucleome Informatics Private Limited, Hyderabad, India.

### Sequencing analysis and gene mapping

Raw reads were obtained from the service provider and processed using *FastQC* to assess the quality. The reads were then used for aligning against the reference genome (Nipponbare) using *BWA mem*. The initial alignment (.*bam*) files were sorted, and duplicates were removed using *samtools.* The analysis-ready bam files were used for calling variants using *freebayes* and the variants were filtered and processed using *VCFtools* and *bcftools*. The variants were filtered to retain only Single Nucleotide Polymorphisms (SNPs) with a mapping quality greater than 30, a minimum and the maximum depth of 10 and 200, respectively. The filtered SNPs were then provided as input to the R language-based QTL mapping tool called *QTLseqr* with default filtering criteria and 0.5 Mb window size. The *QTLseqr* identifies QTL intervals by calculating two parameters namely, Delta-SNP-index and the G’ statistics (Mansfeld and Grumet, 2018; Magwene *et al*., 2011). The DNA sequence and analysis statistics are provided in Table S1.

### SNP annotation and analyses

Information of the putative QTL intervals obtained from the QTL-seq analysis was used to obtain the variations in the intervals. *Bedtools* (Quinlan and Hall, 2010) was used to extract the SNPs in the QTL interval from the whole genome variant file. The QTL SNPs were then annotated using *Variant Effect Predictor* (VEP) to identify the effect of the SNPs on the genome. Microsoft Excel was used for further analyses like segregating the SNPs based on their effects.

### Marker development and validation

SNPs having a high impact on the associated genes were chosen based on the effect (non-sense mutations) and SIFT score (missense mutations causing intolerable amino acid changes) provided by VEP. The presence and absence of selected SNPs in SM92 (mutant line) and SM (the wildtype line), respectively, were confirmed through Sanger’s sequencing. A total of 50 SNPs (42 from the QTL intervals and 8 from the rest of the genome) were finally selected for developing Kompetitive Allele Specific PCR (KASP^TM^) assays (LGC Genomics, UK). The F_2_ individuals derived from SM92 X SM cross were genotyped using KASP assays for all 50 SNPs in the ABI ViiA7 RT-PCR System or Bio-Rad CFX384 system. The genotyping was performed in a 5 µl reaction per assay under conditions recommended by the manufacturer. KASP reaction condition was as follows: 94 LJ -15 minutes; 10 cycles of 94LJ for 20 seconds and 61-55LJ for 1 minute (touch-down of 0.6LJ/cycle); 26 cycles of 94LJ for 20 seconds and 55LJ for 1 minute. TASSEL was used to statistically assess the association between the genotypes and YSB tolerance.

### RNA sequencing and analysis

SM and SM92 plants were infested with YSB larvae during the tillering stage. Stem samples were collected from the infested stems an area 1 cm on either side of the larval entry hole 3 days after release of larvae. Also, uninfested stem samples were collected as a control. All the samples were collected as duplicates. The samples were flash-frozen immediately after collection and stored at −80 until further processing. RNA was isolated from the samples using RNeasy Plant RNA Isolation Kit (Qiagen, Germany) with on-column DNA digestion using RNase-Free DNase Set (Qiagen, Germany). Purified RNA was quantified using Qubit^TM^ RNA HS Assay Kit (Thermo Fisher Scientific, USA). RNA integrity was assessed using agarose gel electrophoresis and BioAnalyzer 2100 (Agilent Technologies, USA). 1 µg of total RNA samples was used for library preparation for each sample using the Illumina TruSeq Stranded Total RNA with Ribo-Zero Plant (Illumina, USA). Sequencing was performed on Illumina’s NovaSeq6000 platform with cycling conditions to obtain 2x100bp reads on an S2 flowcell to obtain ∼70 million reads per sample. The RNA sequence statistics are provided in Table S2. Library preparation and sequencing were performed in the NGS facility of CSIR-CCMB, Hyderabad.

The raw data was obtained, and the quality was assessed using *FastQC*. Adapters and low-quality reads were trimmed using *Trim Galore!* (Krueger)*. RNA-STAR* was used to align the trimmed reads to the rice genome (MSU Rice Genome Annotation Project - version 7) (Dobin *et al*., 2013). Reads that mapped to the genes were quantified using *featureCounts* (Liao *et al*., 2014). Differential gene expression was obtained using the R library *DEseq2* (Love *et al*., 2014). Genes with a false discovery rate ≤ 0.05 and log_2_ fold change ≤ −2 or ≥ 2 were considered differentially expressed genes (DEGs). For gene ontology, and pathway analyses of the DEGs, tools including gProfiler, ShinyGO, and MapMan were used (Thimm *et al*., 2004). *Venny* was used for generating Venn diagrams (Oliveros, 2007).

### Structure prediction and homology search

Amino acid sequences of the DEGs classified under the Lipid Transport/Metabolism category were retrieved from RAPDB (<u>RAP-DB</u>; Sakai et al., 2013; Kawahara et al., 2013). The sequences were subjected to structure prediction using ColabFold (<u>AlphaFold2.ipynb - Colaboratory</u>) which uses the AlphaFold2 algorithm to predict 3D structure models of proteins (Mirdita *et al*., 2022). In order to identify the structural homologues of the predicted structures, a structural homology search was performed using the PDB search option of DALI server (<u>Dali server</u>; Holm, 2022). Multiple sequence alignment of protein sequences and the phylogenetic tree were generated using <u>Clustal</u> <u>Omega</u>(Sievers *et al*., 2011; Goujon *et al*., 2010). The phylogenetic tree visualization and annotation were performed using <u>iTOL</u> (Letunic and Bork, 2021).

### Metabolite profiling and analysis

SM and SM92 plants were infested with YSB larvae during the tillering stage. Stem samples were collected from the infested stems an area 1 cm on either side of the larval entry hole 3 days after release of larvae. Also, uninfested stem samples were collected as a control. All the samples were collected in triplicates. The samples were flash-frozen immediately after collection and stored at −80 until further processing. Further, the frozen samples were lyophilized thoroughly, weighed, and 30 mg of the lyophilized samples were powdered. Metabolites were extracted in 1ml of 80% methanol at 30°C for 30 minutes in shaking condition before centrifuging the samples at 14000rpm for 5 minutes. The supernatant was filtered through 0.22µm membrane. The extracts (10µl each) were analysed by the Shimadzu Prominence HPLC coupled with Shimadzu triple quadrupole LCMS-8045 mass spectrometer (Shimadzu Corporation, Kyoto, Japan). HPLC was performed with a CLC0181 column (Shimpack Gist C18 75*4.0mm, 3μm) with a runtime of 50 minutes at a flow rate of 0.8mL/minute and column temperature of 30°C. Mobile phases used for HPLC include 0.04% (v/v) acetic acid in water and 100% acetonitrile. The electrospray ionization (ESI)–MS analysis was performed in both positive and negative ion modes. Full-scan mass spectra were acquired over a mass range of m/z 50–2000. The sample extraction and metabolite profiling were outsourced to Novelgene Technologies Pvt. Ltd., Hyderabad, India.

The raw intensity data of the detected metabolites was analysed using the one factor statistical analysis module of MetaboAnalyst 5.0 using KEGG compound identifiers (Pang *et al*., 2021). The data was normalized using quantile normalization method, log-transformed, and scaled using the auto-scaling option. The results were downloaded and further processed in Microsoft Excel.

## DATA AVAILABILITY

SM and SM92 parent whole genome sequences are deposited in NCBI-SRA under the accession PRJNA658718. The F_2_ bulk DNA data are deposited under the accession PRJNA972670. The transcriptome data are deposited in NCBI-Gene Expression Omnibus (GEO) under the accession GSE245213.

## AUTHOR CONTRIBUTIONS

Conceptualisation - RVS, MSM, HKP, and PAP. Investigation - GCG (QTL-seq, RNA-seq, metabolomics analyses and KASP genotyping); UB, VB, SB, PAP (YSB larvae rearing, F_2_ population generation, and screening of rice plants); DR (QTLseq data analysis); PV, NM, KJ, SM, PAP (screening of plants in various seasons). Assessment and supervision of the work - GSL, SRLV, KMB, MSR, PAP, HKP, MSM, and RVS. Writing of original draft – GCG. Manuscript review and editing - PAP, KMB, HKP, MSM, and RVS. Funding acquisition – MSM, HKP, and RVS.

## ACKNOWLEDGEMENTS

This study was supported by grants to MSM, HKP, and RVS from the Council of Scientific and Industrial Research (CSIR), Government of India (MLP0121 Phase-I and Phase-II) and the JC Bose Fellowship of RVS granted by the Science and Engineering Research Board (SERB), Department of Science and Technology, Government of India (GAP0444). We thank Dr. Rajkanwar Nathawat (Current address: Yale University) for her guidance and suggestions on protein structure prediction analysis. We thank the skilled workers at ICAR-IIRR and CSIR-CCMB for their support in field activities and maintenance.

## CONFLICT OF INTEREST

The authors declare that no conflict of interest exists.

### Supporting Information

**Table S1: S**tatistics of the whole genome sequence data generated in this study.

**Table S2: S**tatistics of the RNA sequence data generated in this study.

**Table S3:** Overlapping insect resistance QTL intervals

**Table S4:** Laccase-encoding genes that are situated in Chromosome 1 QTL interval.

**Figure S1:** Functional profiling of genes present in the QTL intervals and carrying SNPs.

**Figure S2:** Differential expression of genes in SM and SM92.\

**Figure S3:** Differential regulation of phenylpropanoid pathway genes.

**Figure S4:** Enrichment of lignin metabolism related genes in the QTL intervals.

**Figure S5:** Differentially expressed lipid-related genes and their phylogeny.

**Figure S6:** Other significantly enriched pathways and corresponding differentially expressed genes.

**Figure S7:** Variation in the regulatory region of *OsLTPL146* in SM and SM92.

**Figure S8:** Multiple phenylpropanoid pathway metabolites are accumulated in SM and SM92.

**Data S1:** Phenotype scores of F2 progenies

**Data S2:** List of genes present in all the identified QTL intervals, the SNPs and their effects.

**Data S3:** List of genes predicted to harbor high effect causing SNPs in the QTL regions.

**Data S4:** List of SNPs selected as markers for trait association study using KASP assays.

**Data S5:** KASP assay information

**Data S6:** Genotype-phenotype information of screened F2 progenies

**Data S7:** Marker-Trait Association Analysis of F2 progenies

**Data S8:** Differentially expressed genes in SM upon YSB infestation at 72hpi (infested vs uninfested)

**Data S9:** Differentially expressed genes in SM92 upon YSB infestation at 72hpi (infested vs uninfested)

**Data S10:** List of differentially expressed genes that are located within the QTL intervals and their log2 fold change values.

**Data S11:** Log2 fold change values of genes upregulated upon striped stem borer and yellow stem borer in respective tolerant and susceptible lines

**Data S12:** Raw peak intensity data of metabolites

**Data S13:** Normalized peak intensity data of metabolites

## References

Bentur JS, Padmakumari AP, Jhansilakshmi V, Padmavathi Ch, Kondala Rao Y, A.S. and P.I. (2011) Insect resistance in Rice (2000-2009).,.

Dobin, A., Davis, C.A., Schlesinger, F., Drenkow, J., Zaleski, C., Jha, S., Batut, P., Chaisson, M. and Gingeras, T.R. (2013) STAR: ultrafast universal RNA-seq aligner. Bioinformatics, 29, 15–21. Available at: https://academic.oup.com/bioinformatics/article/29/1/15/272537 [Accessed April 23, 2023].

Dong, N.Q. and Lin, H.X. (2021) Contribution of phenylpropanoid metabolism to plant development and plant–environment interactions. J. Integr. Plant Biol., 63, 180–209. Available at: https://onlinelibrary.wiley.com/doi/full/10.1111/jipb.13054 [Accessed April 23, 2023].

Fauth, M., Schweizer, P., Buchala, A., Markstädter, C., Riederer, M., Kato, T. and Kauss, H. (1998) Cutin Monomers and Surface Wax Constituents Elicit H2O2 in Conditioned Cucumber Hypocotyl Segments and Enhance the Activity of Other H2O2 Elicitors. Plant Physiol., 117, 1373. Available at: /pmc/articles/PMC34901/ [Accessed October 14, 2022].

Franco, O.L., Rigden, D.J., Melo, F.R. and Grossi-de-Sá, M.F. (2002) Plant α-amylase inhibitors and their interaction with insect α-amylases. Eur. J. Biochem., 269, 397–412. Available at: https://onlinelibrary.wiley.com/doi/full/10.1046/j.0014-2956.2001.02656.x [Accessed October 14, 2022].

Gangwar, S.K., Chakraborty, S., Dasgupta, M.K. and Huda, A.K.S. (1986) MODELLING YIELD LOSS IN INDICA RICE IN FARMERS’ FIELDS DUE TO MULTIPLE PESTS. Ecosyst. Environ., 17, 165–171.

Gao, H., Ma, K., Ji, G., Pan, L. and Zhou, Q. (2022) Lipid transfer proteins involved in plant– pathogen interactions and their molecular mechanisms. Mol. Plant Pathol. Available at: https://onlinelibrary.wiley.com/doi/full/10.1111/mpp.13264 [Accessed October 13, 2022].

Gesteiro, N., Butrón, A., Estévez, S. and Santiago, R. (2021) Unraveling the role of maize (Zea mays L.) cell-wall phenylpropanoids in stem-borer resistance. Phytochemistry, 185, 112683.

Goujon, M., McWilliam, H., Li, W., Valentin, F., Squizzato, S., Paern, J. and Lopez, R. (2010) A new bioinformatics analysis tools framework at EMBL–EBI. Nucleic Acids Res., 38, W695– W699. Available at: https://academic.oup.com/nar/article/38/suppl_2/W695/1097251 [Accessed August 12, 2022].

Gururaj Katti, Chitra Shanker, Padmakumari A.P., P.I.C. (2011) Rice stem borers in India-species composition and distribution. Tech. Bull. No. 59., 89.

Heinrichs, Elvis & Medrano, F.G. and Rapusas, H.R. (1985) Genetic Evaluation for Insect Resistance in Rice., Available at: http://books.irri.org/9711041103_content.Pdf.

Holm, L. (2022) Dali server: structural unification of protein families. Nucleic Acids Res., 50, W210–W215. Available at: https://academic.oup.com/nar/article/50/W1/W210/6591528 [Accessed October 13, 2022].

IRRI (2014) *Standard evaluation system for rice. International Rice Research Institute*, Available at: http://www.clrri.org/ver2/uploads/SES_5th_edition.pdf [Accessed July 31, 2020].

Jena, K.K., Jeung, J.U., Lee, J.H., Choi, H.C. and Brar, D.S. (2006) High-resolution mapping of a new brown planthopper (BPH) resistance gene, Bph18(t), and marker-assisted selection for BPH resistance in rice (Oryza sativa L.). Theor. Appl. Genet., 112, 288–297. Available at: https://link.springer.com/article/10.1007/s00122-005-0127-8 [Accessed July 1, 2022].

Jumper, J., Evans, R., Pritzel, A., et al. (2021) Highly accurate protein structure prediction with AlphaFold. Nat. 2021 5967873, 596, 583–589. Available at: https://www.nature.com/articles/s41586-021-03819-2 [Accessed October 14, 2022].

Kawahara, Y., la Bastide, M. de, Hamilton, J.P., et al. (2013) Improvement of the oryza sativa nipponbare reference genome using next generation sequence and optical map data. Rice, 6, 3–10. Available at: https://thericejournal.springeropen.com/articles/10.1186/1939-8433-6-4 [Accessed August 12, 2022].

Kim, T.H., Park, J.H., Kim, M.C. and Cho, S.H. (2008) Cutin monomer induces expression of the rice OsLTP5 lipid transfer protein gene. J. Plant Physiol., 165, 345–349. Available at: https://linkinghub.elsevier.com/retrieve/pii/S0176161707001162 [Accessed August 11, 2020].

Kolattukudy, P.E., Rogers, L.M., Li, D., Hwang, C.S. and Flaishman, M.A. (1995) Surface signaling in pathogenesis. Proc. Natl. Acad. Sci. U. S. A., 92, 4080. Available at: /pmc/articles/PMC41890/?report=abstract [Accessed October 14, 2022].

Krueger, F. TrimGalore: A wrapper around Cutadapt and FastQC to consistently apply adapter and quality trimming to FastQ files, with extra functionality for RRBS data. Available at: https://github.com/FelixKrueger/TrimGalore [Accessed April 23, 2023].

Lage, J.-L. Da (2018) The Amylases of Insects. Int. J. Insect Sci., 10, 117954331880478. Available at: /pmc/articles/PMC6176531/ [Accessed October 14, 2022].

Lee, M.-H., Jeon, H.S., Kim, S.H., et al. (2019) Lignin-based barrier restricts pathogens to the infection site and confers resistance in plants. EMBO J., 38, e101948. Available at: https://onlinelibrary.wiley.com/doi/full/10.15252/embj.2019101948 [Accessed October 13, 2022].

Letunic, I. and Bork, P. (2021) Interactive Tree Of Life (iTOL) v5: an online tool for phylogenetic tree display and annotation. Nucleic Acids Res., 49, W293–W296. Available at: https://academic.oup.com/nar/article/49/W1/W293/6246398 [Accessed August 12, 2022].

Liao, Y., Smyth, G.K. and Shi, W. (2014) featureCounts: an efficient general purpose program for assigning sequence reads to genomic features. Bioinformatics, 30, 923–930. Available at: https://academic.oup.com/bioinformatics/article/30/7/923/232889 [Accessed April 23, 2023].

Love, M.I., Huber, W. and Anders, S. (2014) Moderated estimation of fold change and dispersion for RNA-seq data with DESeq2. Genome Biol., 15, 1–21. Available at: https://genomebiology.biomedcentral.com/articles/10.1186/s13059-014-0550-8 [Accessed April 23, 2023].

M.D. Pathak and Z.R. Khan (1994) *Insect Pests of Rice*, International Rice Research Institute. Available at: http://books.irri.org/9712200280_content.pdf [Accessed December 6, 2021].

Magwene, P.M., Willis, J.H. and Kelly, J.K. (2011) The Statistics of Bulk Segregant Analysis Using Next Generation Sequencing. PLOS Comput. Biol., 7, e1002255. Available at: https://journals.plos.org/ploscompbiol/article?id=10.1371/journal.pcbi.1002255 [Accessed October 23, 2022].

Malvar, R.A., Butrón, A., Ordás, B. and Santiago, R. (2008) Causes of natural resistance to stem borers in maize. Crop Prot. Res. Adv., 313.

Mansfeld, B.N. and Grumet, R. (2018) QTLseqr: An R Package for Bulk Segregant Analysis with NextLJGeneration Sequencing. Plant Genome, 11, 180006. Available at: https://onlinelibrary.wiley.com/doi/10.3835/plantgenome2018.01.0006 [Accessed December 25, 2020].

Miller, G.L. (1959) Use of Dinitrosalicylic Acid Reagent for Determination of Reducing Sugar. Anal. Chem., 31, 426–428. Available at: https://pubs.acs.org/doi/abs/10.1021/ac60147a030 [Accessed August 29, 2023].

Mirdita, M., Schütze, K., Moriwaki, Y., Heo, L., Ovchinnikov, S. and Steinegger, M. (2022) ColabFold: making protein folding accessible to all. Nat. Methods 2022 196, 19, 679–682. Available at: https://www.nature.com/articles/s41592-022-01488-1 [Accessed August 12, 2022].

Moin, M., Bakshi, A., Saha, A., Dutta, M., Madhav, S.M. and Kirti, P.B. (2016) Rice ribosomal protein large subunit genes and their spatio-temporal and stress regulation. Front. Plant Sci., 7, 1284.

Muralidharan, K. and Pasalu, I.C. (2006) Assessments of crop losses in rice ecosystems due to stem borer damage (Lepidoptera: Pyralidae). Crop Prot., 25, 409417.

Nagaraj, S., Senthil-Kumar, M., Ramu, V.S., Wang, K. and Mysore, K.S. (2016) Plant ribosomal proteins, RPL12 and RPL19, play a role in nonhost disease resistance against bacterial pathogens. Front. Plant Sci., 6, 1192.

Oliveros, J.C. (2007) Venny. An interactive tool for comparing lists with Venn’s diagrams. Available at: https://bioinfogp.cnb.csic.es/tools/venny/index.html.

Oneda, H., Lee, S. and Inouye, K. (2004) Inhibitory Effect of 0.19 α-Amylase Inhibitor from Wheat Kernel on the Activity of Porcine Pancreas α-Amylase and Its Thermal Stability. J. Biochem., 135, 421–427. Available at: https://academic.oup.com/jb/article/135/3/421/1072114 [Accessed October 14, 2022].

Padmakumari A.P, Seshu Madhav M, Uma Kanth B, Sunil B, Gopi, P, Karteek J, Laha GS, Sundaram, RM, Subba Rao, LV, H.K.P. and R.S. (2017) EMS mutants as a novel source of tolerance to yellow stem borer., Hyderabad.

Padmakumari AP and Katti G (2018) Designing insect bioassays for studying components of insect pest management-rice yellow stem borer – as a case study. In National Symposium Entomology2018:Advances and Challenges. Hyderabad, p. 117.

Pang, Z., Chong, J., Zhou, G., et al. (2021) MetaboAnalyst 5.0: narrowing the gap between raw spectra and functional insights. Nucleic Acids Res., 49, W388–W396. Available at: https://academic.oup.com/nar/article/49/W1/W388/6279832 [Accessed April 23, 2023].

Pathak, M.D. (1977) DEFENSE OF THE RICE CROP AGAINST INSECT PESTS. Ann. N. Y. Acad. Sci., 287, 287–295. Available at: https://onlinelibrary.wiley.com/doi/full/10.1111/j.1749-6632.1977.tb34247.x [Accessed December 6, 2021].

Pathak, M.D. (1968) Ecology of Common Insect Pests of Rice. 10.1146/annurev.en.13.010168.001353, 13, 257–294. Available at: https://www.annualreviews.org/doi/abs/10.1146/annurev.en.13.010168.001353 [Accessed December 6, 2021].

Peltier, A.J., Hatfield, R.D. and Grau, C.R. (2009) Soybean Stem Lignin Concentration Relates to Resistance to Sclerotinia sclerotiorum. Plant Dis., 93, 149–154. Available at: https://pubmed.ncbi.nlm.nih.gov/30764097/ [Accessed October 13, 2022].

Potupureddi, G., Balija, V., Ballichatla, S., et al. (2021) Mutation resource of Samba Mahsuri revealed the presence of high extent of variations among key traits for rice improvement. PLoS One, 16, e0258816. Available at: https://journals.plos.org/plosone/article?id=10.1371/journal.pone.0258816 [Accessed May 24, 2022].

Quinlan, A.R. and Hall, I.M. (2010) BEDTools: a flexible suite of utilities for comparing genomic features. Bioinformatics, 26, 841–842. Available at: https://academic.oup.com/bioinformatics/article/26/6/841/244688 [Accessed May 8, 2023].

Ramu, V.S., Dawane, A., Lee, S., Oh, S., Lee, H.K., Sun, L., Senthil-Kumar, M. and Mysore, K.S. (2020) Ribosomal protein QM/RPL10 positively regulates defence and protein translation mechanisms during nonhost disease resistance. Mol. Plant Pathol., 21, 1481–1494. Available at: https://onlinelibrary.wiley.com/doi/full/10.1111/mpp.12991 [Accessed October 14, 2022].

Rubia, E.G., Heong, K.L., Zalucki, M., Gonzales, B. and Norton, G.A. (1996) Mechanisms of compensation of rice plants to yellow stem borer Scirpophaga incertulas (Walker) injury. Crop Prot., 15, 335–340.

Sakai, H., Lee, S.S., Tanaka, T., et al. (2013) Rice Annotation Project Database (RAP-DB): an integrative and interactive database for rice genomics. Plant Cell Physiol., 54. Available at: https://pubmed.ncbi.nlm.nih.gov/23299411/ [Accessed August 12, 2022].

Salminen, T.A., Blomqvist, K. and Edqvist, J. (2016) Lipid transfer proteins: classification, nomenclature, structure, and function. Planta, 244, 971–997. Available at: https://pubmed.ncbi.nlm.nih.gov/27562524/ [Accessed October 14, 2022].

Santiago, R., Barros-Rios, J. and Malvar, R.A. (2013) Impact of Cell Wall Composition on Maize Resistance to Pests and Diseases. Int. J. Mol. Sci. 2013, Vol. 14, Pages 6960-6980, 14, 6960–6980. Available at: https://www.mdpi.com/1422-0067/14/4/6960/htm [Accessed October 13, 2022].

Santos, R.B., Nascimento, R., Coelho, A. V. and Figueiredo, A. (2020) Grapevine–Downy Mildew Rendezvous: Proteome Analysis of the First Hours of an Incompatible Interaction. Plants 2020, Vol. 9, Page 1498, 9, 1498. Available at: https://www.mdpi.com/2223-7747/9/11/1498/htm [Accessed October 14, 2022].

Savary, S., Willocquet, L., Pethybridge, S.J., Esker, P., McRoberts, N. and Nelson, A. (2019) The global burden of pathogens and pests on major food crops. *Nat*. Ecol. Evol., 3, 430–439. Available at: https://www.nature.com/articles/s41559-018-0793-y.

Schweizer, P., Jeanguenat, A., Whitacre, D., Métraux, J.P. and Mösinger, E. (1996) Induction of resistance in barley againstErysiphe graminisf.sp.hordeiby free cutin monomers. Physiol. Mol. Plant Pathol., 49, 103–120.

Selvaraju, K., Shanmugasundaram, P., Mohankumar, S., Asaithambi, M. and Balasaraswathi, R. (2007) Detection of quantitative trait locus for leaffolder (Cnaphalocrocis medinalis (Guenée)) resistance in rice on linkage group 1 based on damage score and flag leaf width. Euphytica, 157, 35–43. Available at: https://link.springer.com/article/10.1007/s10681-007-9394-6 [Accessed July 1, 2022].

Sievers, F., Wilm, A., Dineen, D., et al. (2011) Fast, scalable generation of high-quality protein multiple sequence alignments using Clustal Omega. Mol. Syst. Biol., 7, 539. Available at: https://onlinelibrary.wiley.com/doi/full/10.1038/msb.2011.75 [Accessed August 12, 2022].

Su, J., Zhang, Z., Wu, M. and Gao, C. (2014) Geographic susceptibility of Chilo suppressalis Walker (Lepidoptera: Crambidae), to chlorantraniliprole in China. Pest Manag. Sci., 70, 989– 995. Available at: https://onlinelibrary.wiley.com/doi/full/10.1002/ps.3640 [Accessed October 13, 2022].

Sun, L., Su, C., Wang, C., Zhai, H. and Wan, J. (2005) Mapping of a Major Resistance Gene to the Brown Planthopper in the Rice Cultivar Rathu Heenati. Breed. Sci., 55, 391–396. Available at: http://www. [Accessed July 1, 2022].

Tae, H.K., Moon, C.K., Jong, H.P., Seong, S.H., Byung, R.K., Byoung, Y.M., Mi, C.S. and Sung, H.C. (2006) Differential expression of rice lipid transfer protein gene(LTP) classes in response to abscisic acid, salt, salicylic acid, and the fungal pathogenMagnaporthe grisea. J. Plant Biol. 2006 495, 49, 371–375. Available at: https://link.springer.com/article/10.1007/BF03178814 [Accessed October 14, 2022].

Takagi, H., Abe, A., Yoshida, K., et al. (2013) QTL-seq: Rapid mapping of quantitative trait loci in rice by whole genome resequencing of DNA from two bulked populations. Plant J., 74, 174–183. Available at: https://onlinelibrary.wiley.com/doi/full/10.1111/tpj.12105 [Accessed July 29, 2020].

Thimm, O., Bläsing, O., Gibon, Y., et al. (2004) MAPMAN: a user-driven tool to display genomics data sets onto diagrams of metabolic pathways and other biological processes. Plant J., 37, 914–939. Available at: https://pubmed.ncbi.nlm.nih.gov/14996223/ [Accessed April 23, 2023].

Trautmann, H.S. and Ramsey, K.M. (2022) A Ribosomal Protein Homolog Governs Gene Expression and Virulence in a Bacterial Pathogen. J. Bacteriol. Available at: https://journals.asm.org/doi/10.1128/jb.00268-22 [Accessed October 14, 2022].

Wang, Y., Ju, D., Yang, X., Ma, D. and Wang, X. (2018) Comparative Transcriptome Analysis Between Resistant and Susceptible Rice Cultivars Responding to Striped Stem Borer (SSB), Chilo suppressalis (Walker) Infestation. Front. Physiol., 9, 1717.

Warner, J.R. and McIntosh, K.B. (2009) How Common Are Extraribosomal Functions of Ribosomal Proteins? Mol. Cell, 34, 3–11.

Weisberg, R.A. (2008) Transcription by Moonlight: Structural Basis of an Extraribosomal Activity of Ribosomal Protein S10. Mol. Cell, 32, 747–748.

Wool, I.G. (1996) Extraribosomal functions of ribosomal proteins. Trends Biochem. Sci., 21, 164– 165.

Yu, Y., Xing, Y., Liu, F., Zhang, X., Li, X., Zhang, J. and Sun, X. (2021) The Laccase Gene Family Mediate Multi-Perspective Trade-Offs during Tea Plant (Camellia sinensis) Development and Defense Processes. Int. J. Mol. Sci. 2021, Vol. 22, Page 12554, 22, 12554. Available at: https://www.mdpi.com/1422-0067/22/22/12554/htm [Accessed October 13, 2022].

Zhao, Q., Nakashima, J., Chen, F., et al. (2013) LACCASE Is Necessary and Nonredundant with PEROXIDASE for Lignin Polymerization during Vascular Development in Arabidopsis. Plant ed October 13, 2022].

