## Supplementary material for "Multiomics-assisted characterization of rice-Yellow Stem Borer interaction provides genomic and mechanistic insights into stem borer tolerance in rice": Figure S

**Supplementary Figures**

**Figure S1 to S8**

**Multiomics-assisted characterization of rice-Yellow Stem Borer interaction provides genomic and mechanistic insights into stem borer tolerance in rice**

Gokulan C. G.^1^, Umakanth Bangale^2^, Vishalakshi Balija^2^, Suneel Ballichatla^2^, Gopi Potupureddi^2^, Deepti Rao^1^, Prashanth Varma^2^, Nakul Magar^2^, Karteek J.^2^, Sravan M.^2^, Padmakumari A. P.^2^, Gouri S Laha^2^, Subba Rao LV^2^, Kalyani M. Barbadikar^2^, Meenakshi Sundaram Raman^2^, Hitendra K. Patel^1,3^, M Sheshu Madhav^2,4, *^, Ramesh V. Sonti^1,^^5,*^

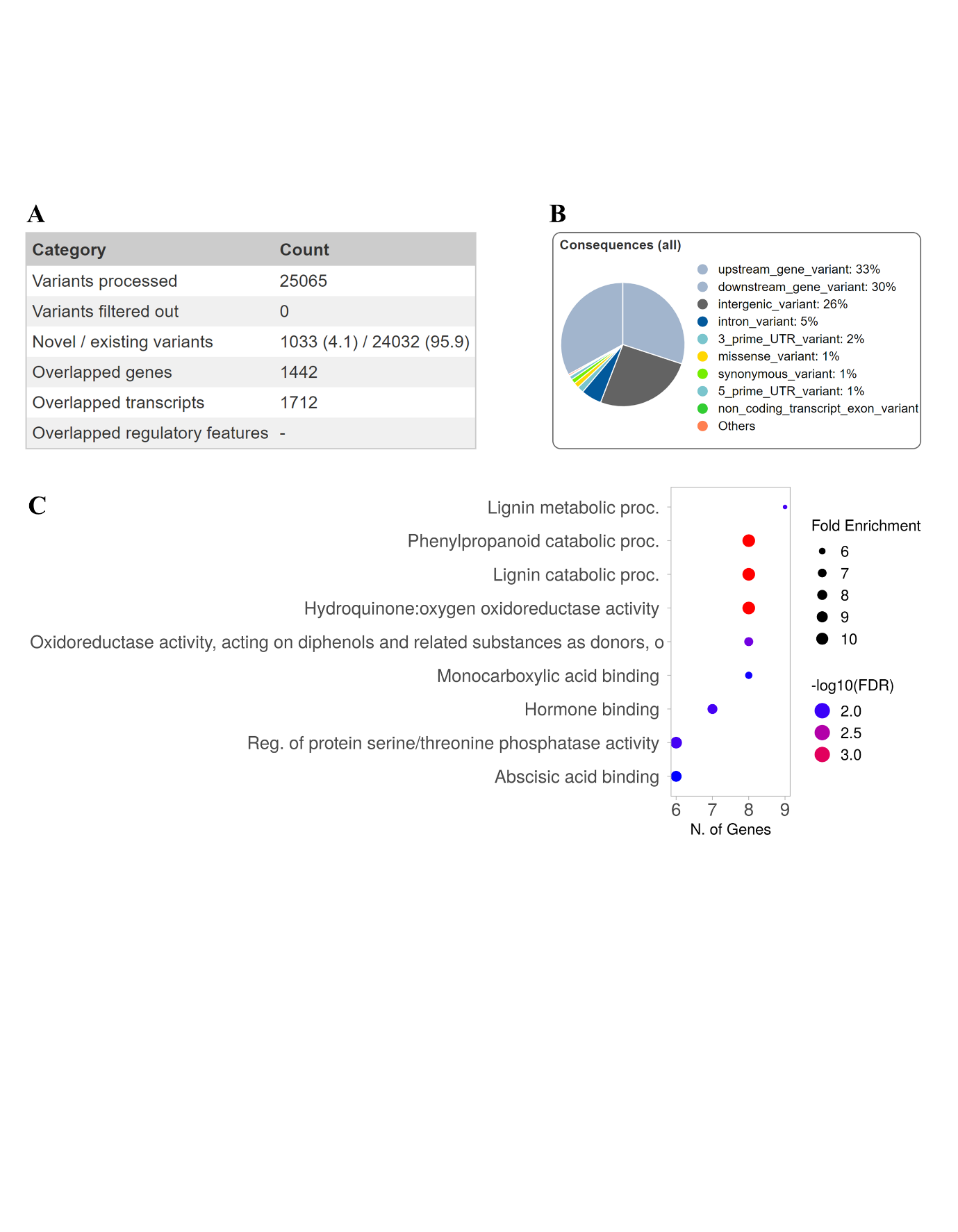

**Figure S1**

Functional profiling of genes present in the QTL intervals and carrying SNPs. (A) Statistical analysis of the variations present within the QTL intervals. (B) Prediction of SNPs effects based on their location in the genome. (C) Enrichment analysis of genes carrying SNPs and located within the QTL intervals showed the presence of multiple genes that are associated with lignin metabolism.

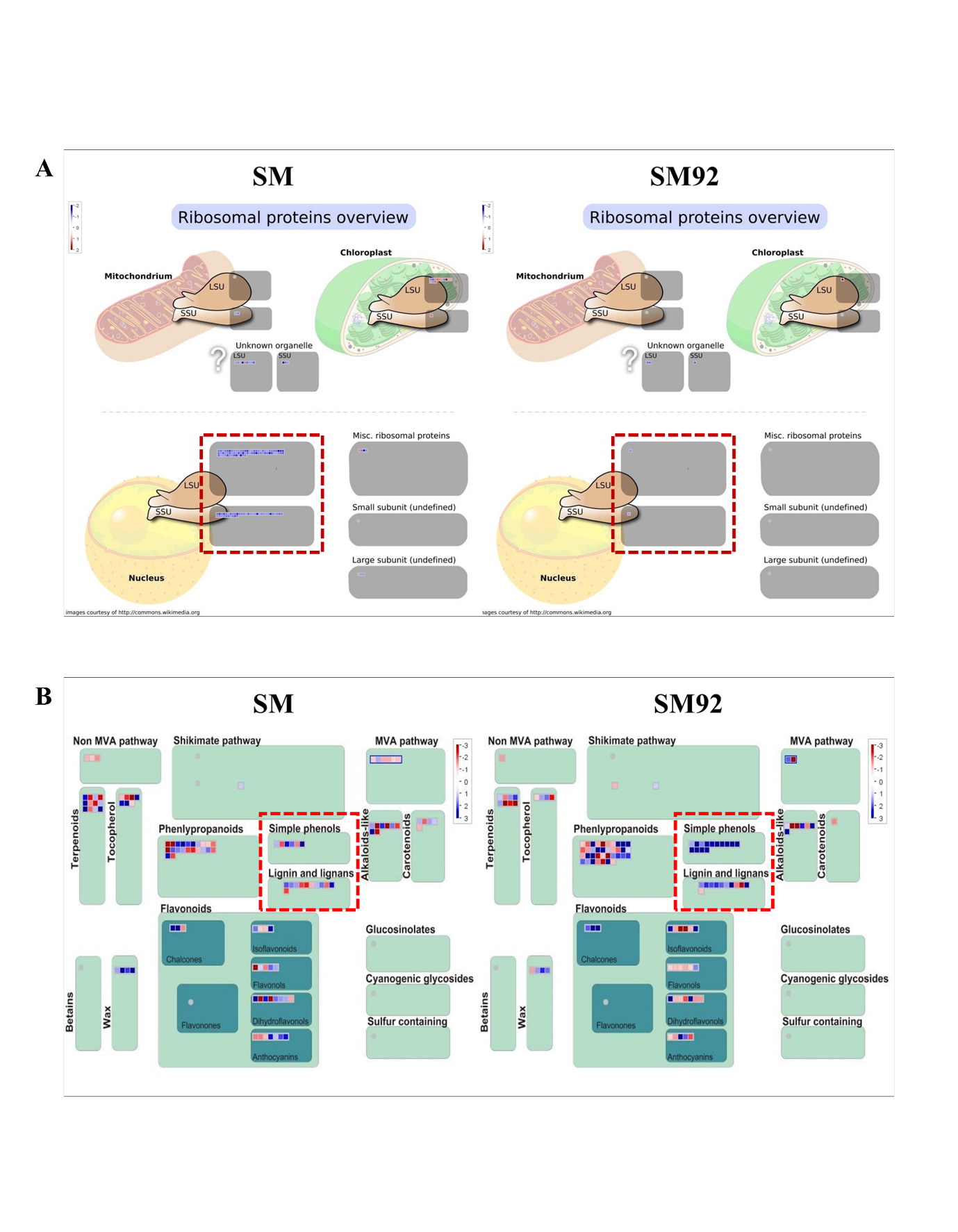

**Figure S2**

Differential expression of genes in SM and SM92. (A) Gene encoding ribosomal subunit proteins (dotted red box) showed exclusive upregulation in SM upon YSB infestation. (B) Multiple genes belonging to simple phenol pathway (red dotted box) were majorly upregulated in SM92 while genes in SM showed mixed regulation upon YSB infestation.

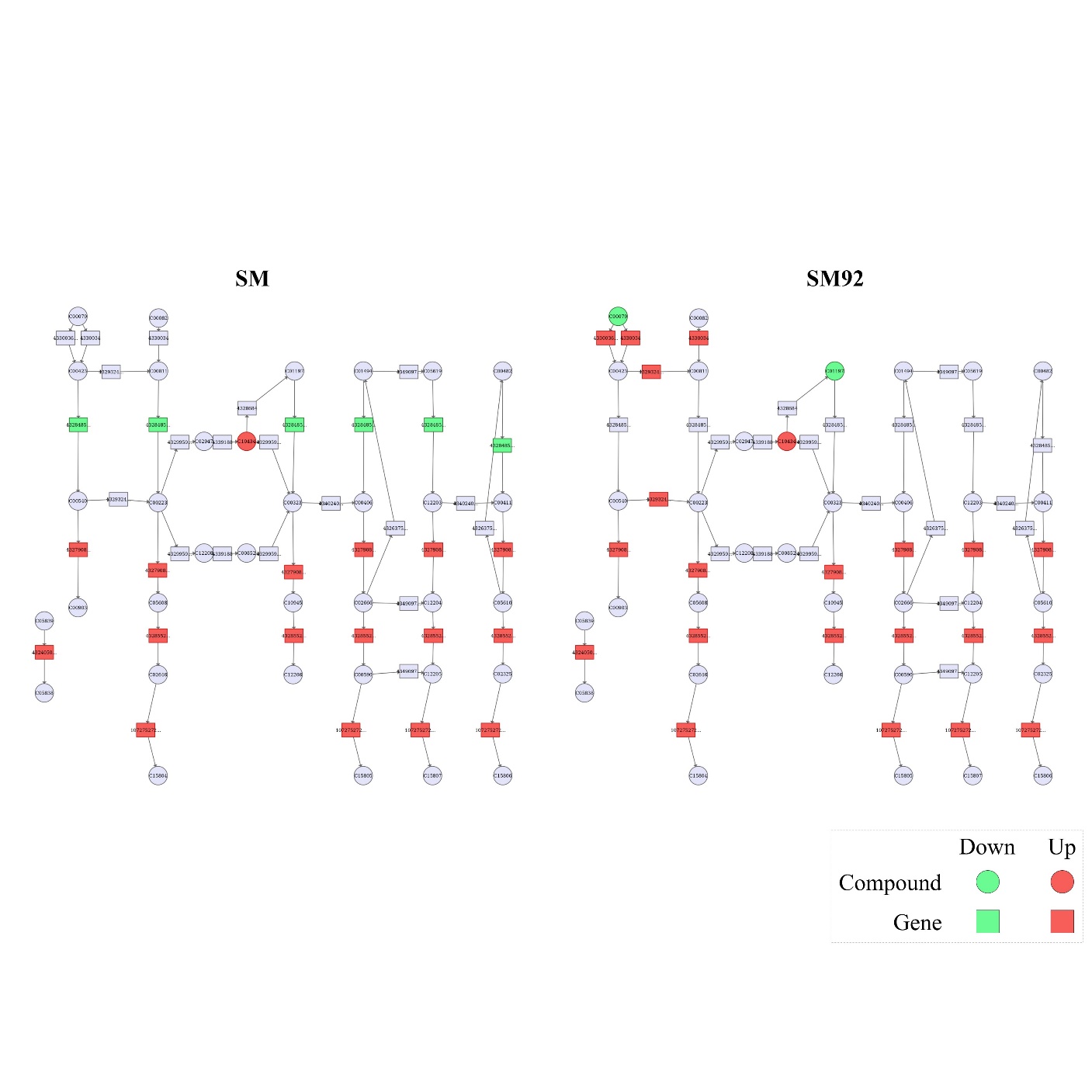

**Figure S3**

Differential regulation of phenylpropanoid pathway genes. Multiple genes belonging to the phenylpropanoid pathway were differentially expressed in SM and SM92 upon YSB infestation. The rectangles represent genes, and the circles represent the compounds. All the differentially expressed genes were upregulated in SM92 (red), whereas in SM, genes encoding proteins catalyzing the early steps in the pathway were downregulated (green).

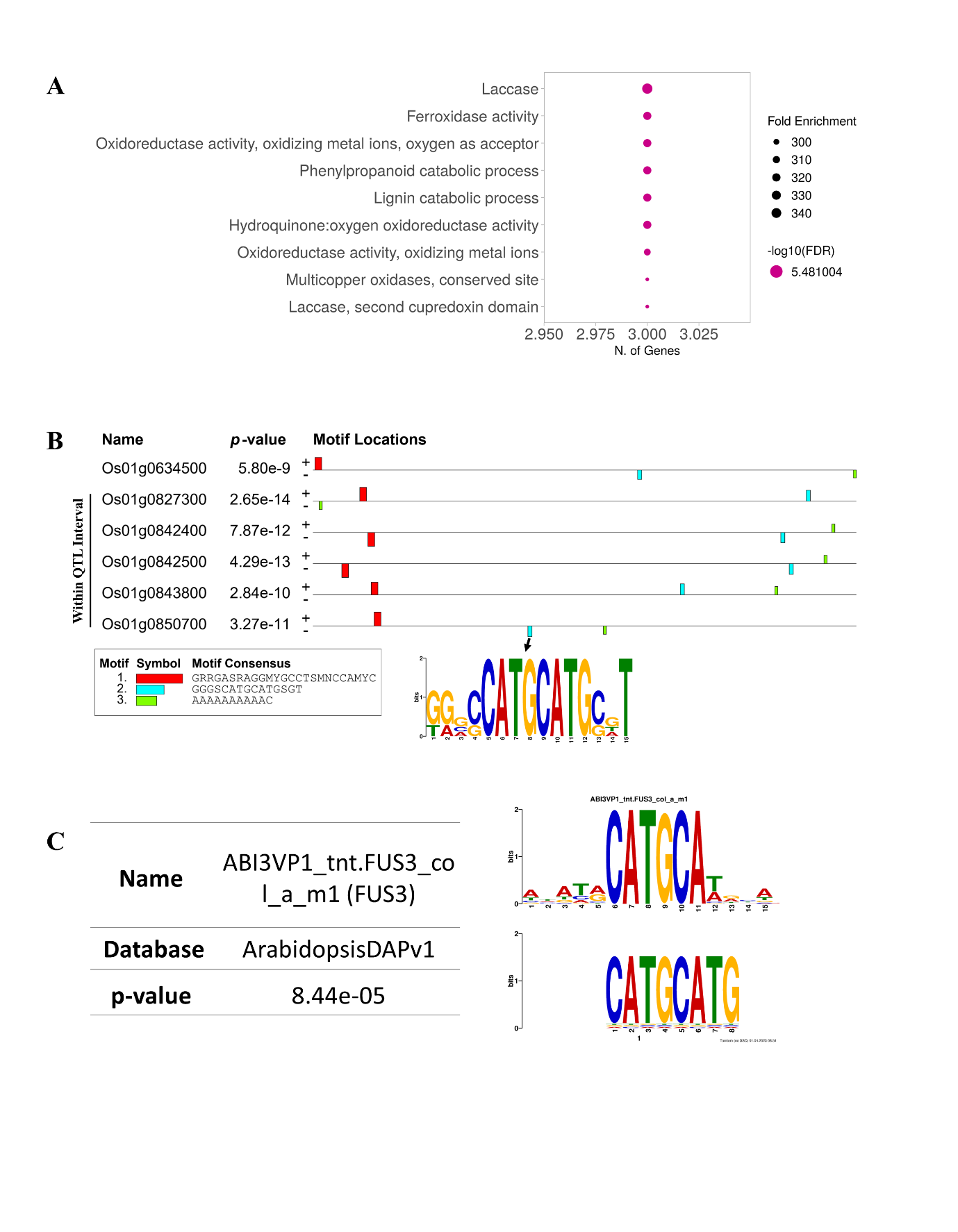

**Figure S4**

Enrichment of lignin metabolism related genes in the QTL intervals. (A) Enrichment analysis of genes from the QTL intervals indicating the enrichment of laccase-encoding genes, that are involved in lignin metabolism. (B) Motif prediction revealed the presence of common motif signatures in the promoters (1kb upstream) of six laccase encoding genes of which are present within the QTL limits. Black arrow denotes the predicted motif 2, which consists of a palindromic motif CATG, present in tandem. (C) Transcription factor prediction analysis indicates that the CATG repeat motif could be a binding site for a B3-domain containing transcription factor using *Arabidopsis* *thaliana* database.

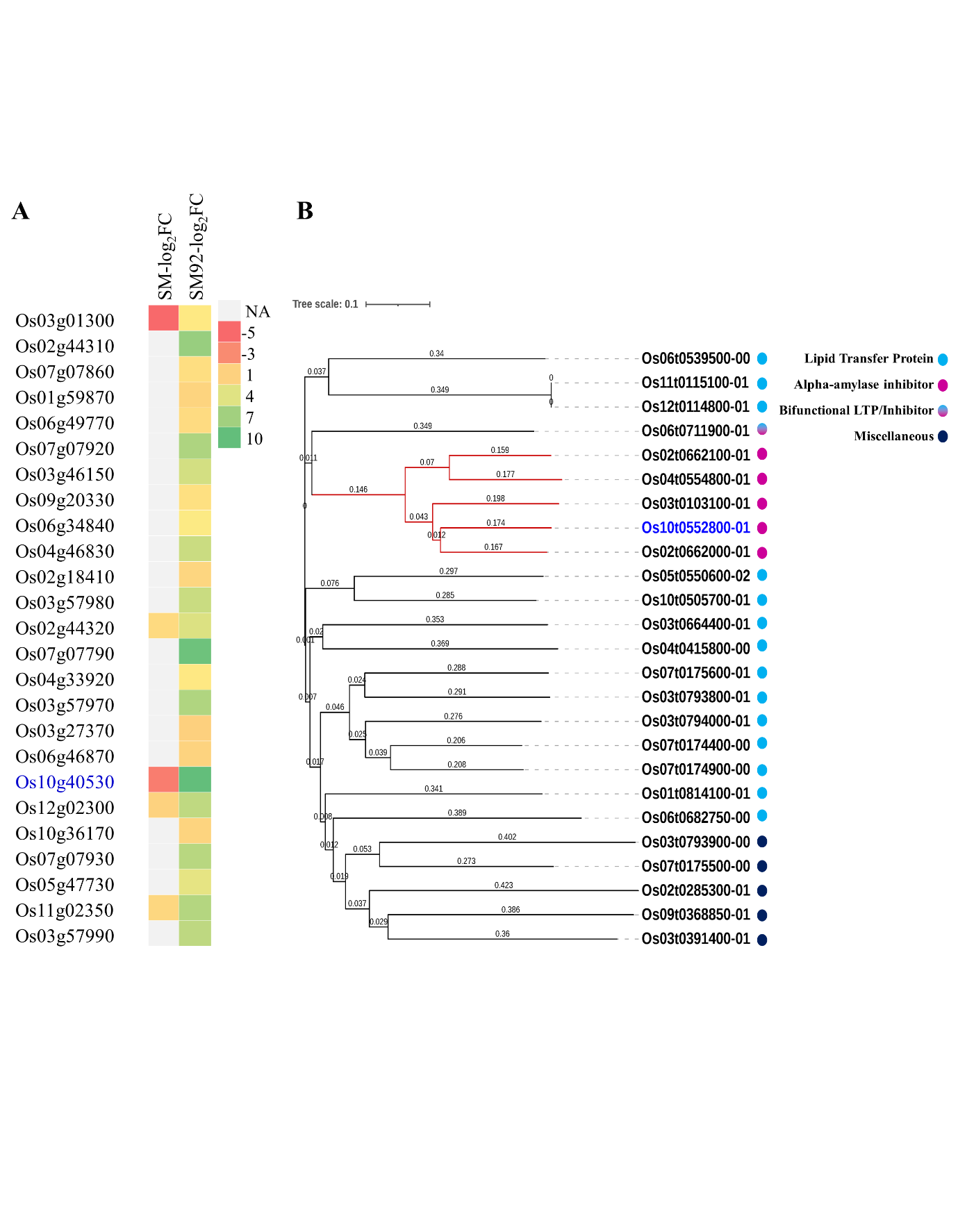

**Figure S5**

Differentially expressed lipid-related genes and their phylogeny. (A) Heatmap showing the log2 fold change values of genes annotated under lipid transfer-like or lipid metabolism related categories in SM and SM92 upon YSB infestation. (B) Phylogenetic tree derived using the multiple sequence alignment of the protein sequences of the genes given in (A). The colored circles beside gene identifiers indicate their putative function as predicted through structure-based analysis. Blue font in (A) and (B) denote *OsLTPL146.* Legends show the log2 fold change value in (A) and the putative function of the proteins in (B). NA indicates not available (no differential expression).

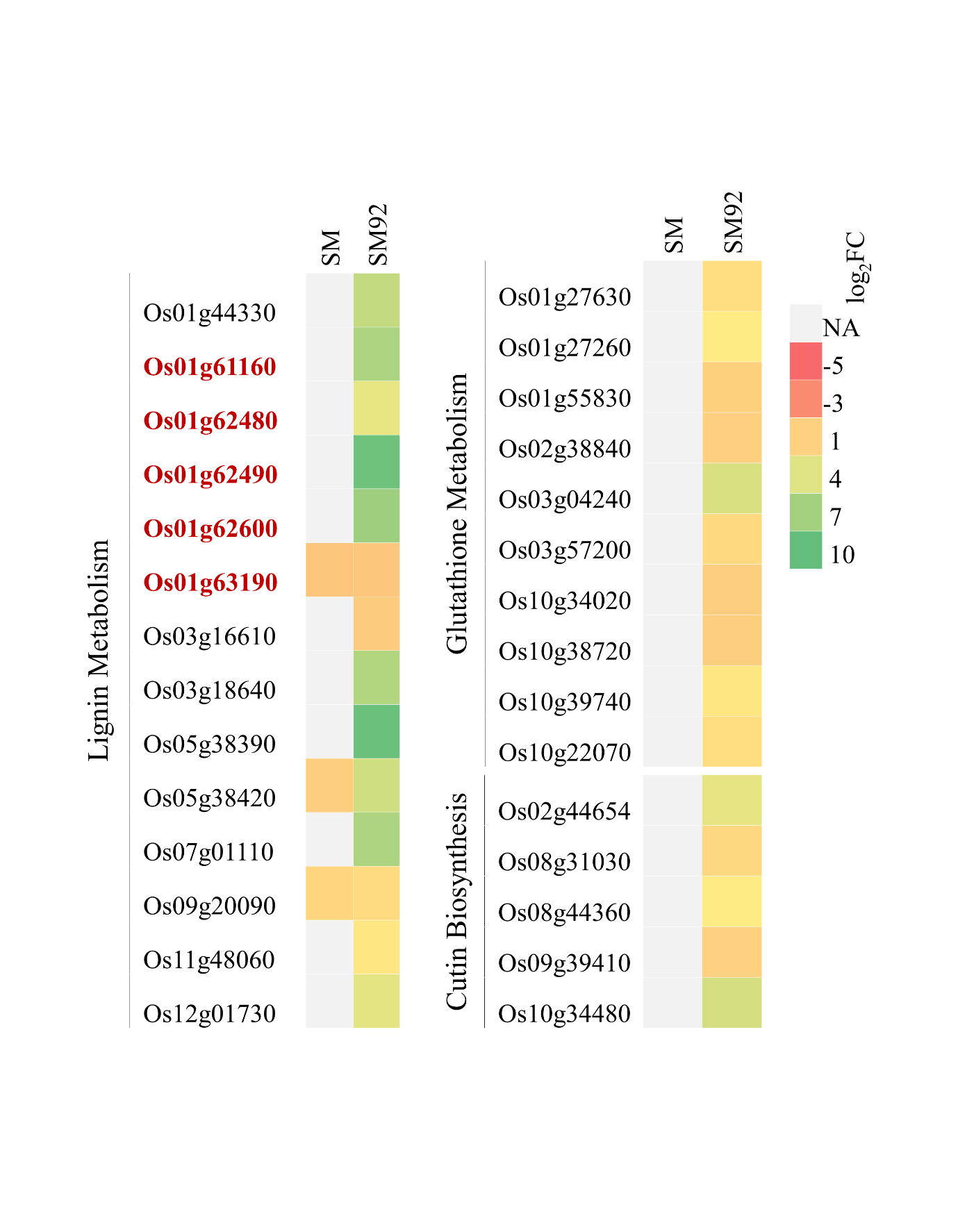

**Figure S6**

Other significantly enriched pathways and corresponding differentially expressed genes. Heatmap showing the log2 fold change values of various genes belonging to multiple pathways including lignin metabolism, glutathione metabolism, and cutin biosynthesis in SM and SM92 upon YSB infestation. Red font gene identifiers indicate the genes located within the QTL intervals. Legends show the log2 fold change. NA indicates not available (no differential expression).

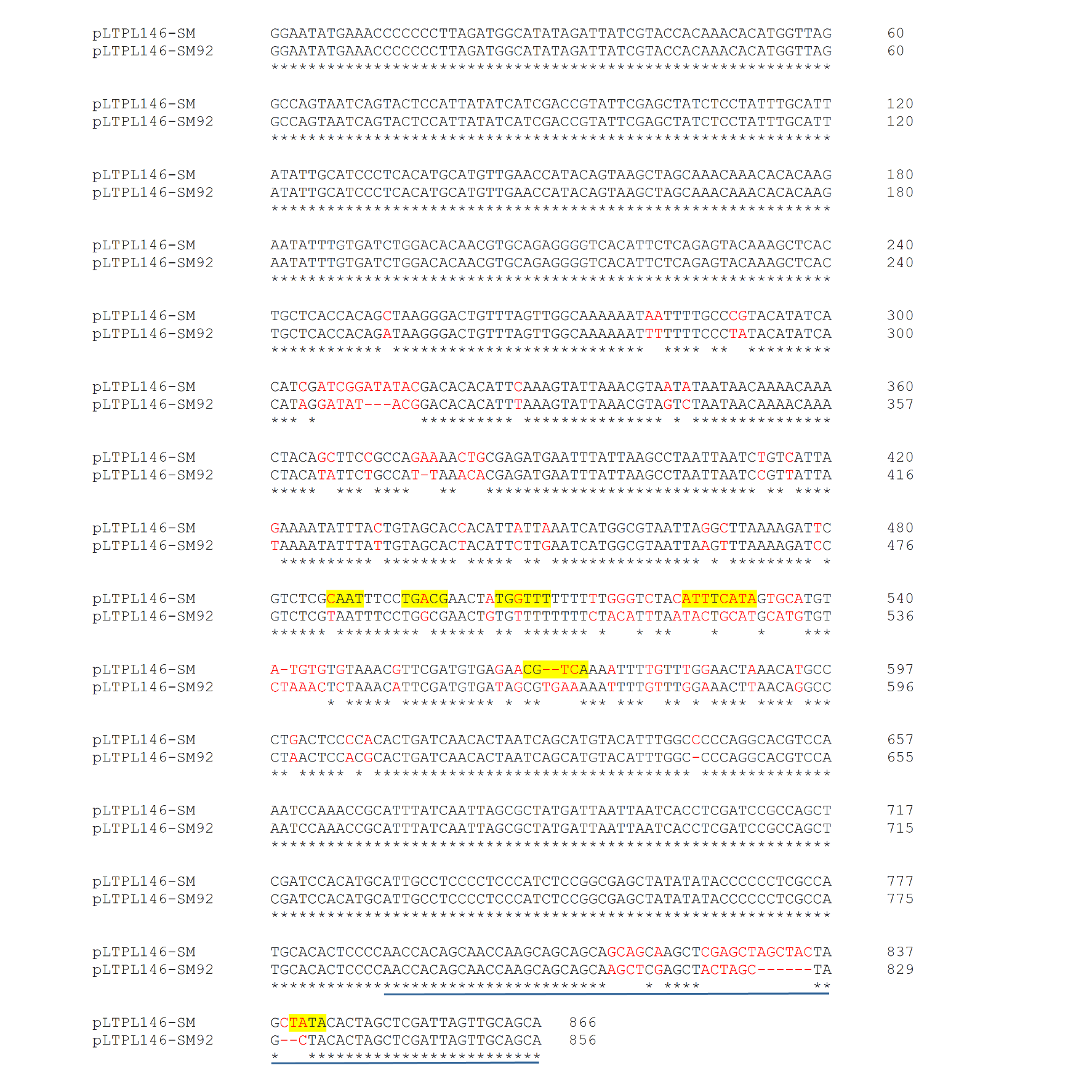
**Figure S7**

Variation in the regulatory region of *OsLTPL146* in SM and SM92. Pairwise alignment of the promoter region of *OsLTPL146* sequenced from SM and SM92 performed using ClustalW. Red fonts indicate bases that vary between SM and SM92. The sequences highlighted in yellow are predicted cis elements in SM which carry variations in SM92. Sequences underlined in blue correspond to the 5 prime untranslated region (5‘ UTR) of *OsLTPL146*.

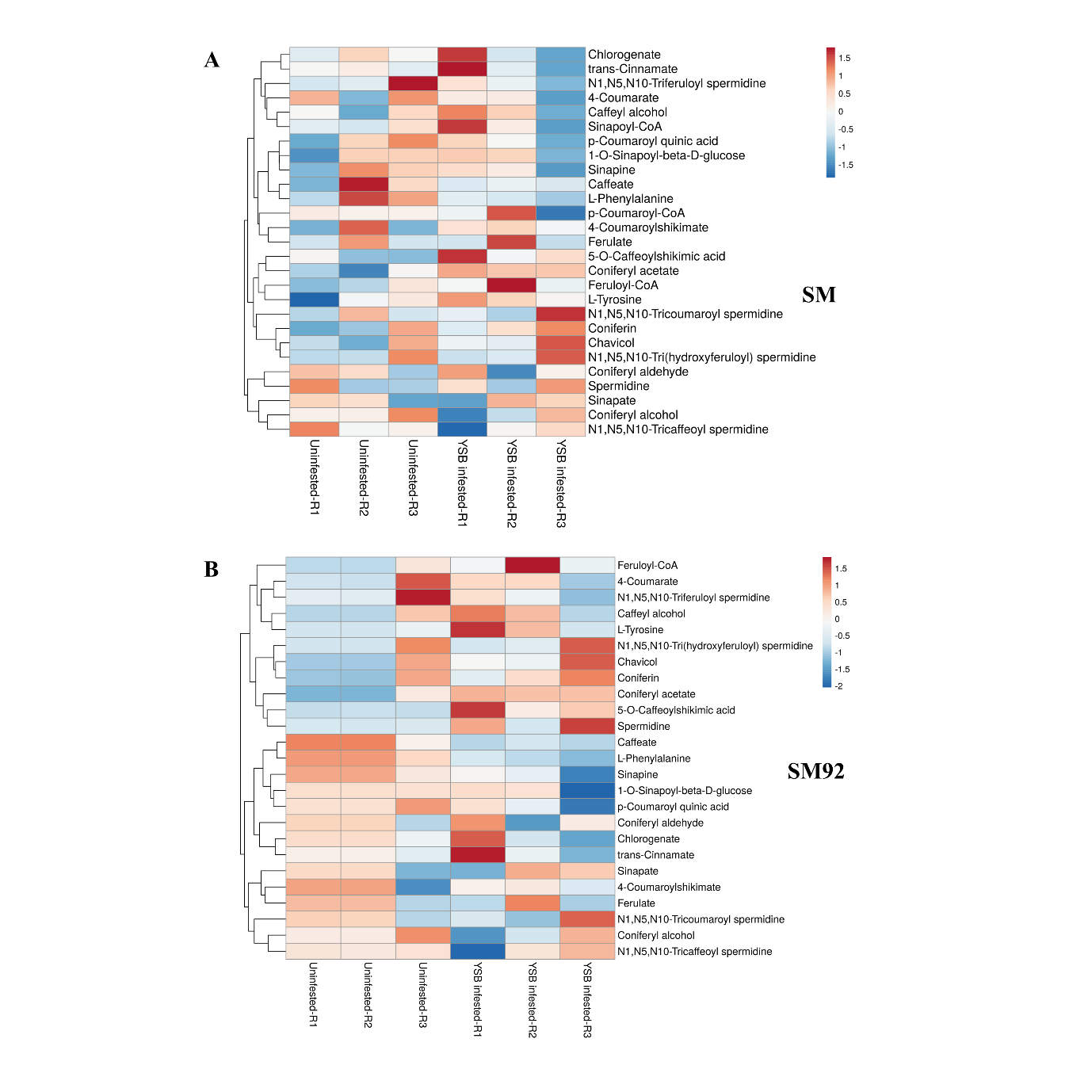

**Figure S8**

Multiple phenylpropanoid pathway metabolites are accumulated in SM and SM92. Heatmap showing the log-transformed signal intensity of metabolites associated with phenylpropanoid pathway in uninfested (Control) and YSB infested conditions in (A) SM and (B) SM92. Legend indicates the range of log-transformed signal intensities.
