## Supplementary material for "Multiomics-assisted characterization of rice-Yellow Stem Borer interaction provides genomic and mechanistic insights into stem borer tolerance in rice": Table S

**Supplementary Tables**

**Table S1 to S4**

**Multiomics-assisted characterization of rice-Yellow Stem Borer interaction provides genomic and mechanistic insights into stem borer tolerance in rice**

Gokulan C. G.^1^, Umakanth Bangale^2^, Vishalakshi Balija^2^, Suneel Ballichatla^2^, Gopi Potupureddi^2^, Deepti Rao^1^, Prashanth Varma^2^, Nakul Magar^2^, Karteek J.^2^, Sravan M.^2^, Padmakumari A. P.^2^, Gouri S Laha^2^, Subba Rao LV^2^, Kalyani M. Barbadikar^2^, Meenakshi Sundaram Raman^2^, Hitendra K. Patel^1,3^, M Sheshu Madhav^2,4, *^, Ramesh V. Sonti^1,^^5,*^

**Table S1: S**tatistics of the whole genome sequence data generated in this study.

| **Samples** | **Raw reads** | **QC passed reads** | **Mapped reads ^a^** | **No. of SNPs ^a^** |
| --- | --- | --- | --- | --- |
| SM | 110937222 | 101706696 | 99665043 | 2637491 |
| SM92 | 108091210 | 97544551 | 95446349 | 2518135 |
| Tolerant Bulk | 120326090 | 118853810 | 116133682 | 2929248 |
| Susceptible Bulk | 112202294 | 110726832 | 108018662 | 2896663 |

^a^ With respect to Nipponbare reference genome (MSU version 7)

**Table S2: S**tatistics of the RNA sequence data generated in this study.

| **Sample** | **Replicate** | **Total reads (x2)** | **Aligned reads ^a^** | **Assigned reads ^a^** |
| --- | --- | --- | --- | --- |
| SM – Uninfested | 1 | 71690714 | 68039389 | 53628824 |
|  | 2 | 82840640 | 78077626 | 54973912 |
| SM92 – Uninfested | 1 | 77130249 | 72224337 | 52315868 |
|  | 2 | 78649531 | 73934930 | 53268945 |
| SM – YSB infested | 1 | 77494218 | 73448373 | 57603629 |
|  | 2 | 73067689 | 68914219 | 54058916 |
| SM92 – YSB infested | 1 | 79116030 | 74023774 | 50540724 |
|  | 2 | 71360146 | 67519603 | 50953185 |

^a^ With respect to Nipponbare reference genome (MSU version 7)

**Table S3:** List of the QTL intervals that overlap between YSB tolerance (this study) and previously reported insect resistance QTL recorded in the QTARO database. Mbp – Mega basepairs.

| **Chrom** | **Insect resistance**  **QTL Database** | | **YSB tolerance QTL intervals** ^a^ | | **Overlap (Mb)** | **Known for resistance to** | **Reference** |
| --- | --- | --- | --- | --- | --- | --- | --- |
|  | **Start** | **End** | **Start** | **End** |  |  |  |
| **1** | 34.5 Mb | 39.6 Mb | 33.4 Mb | 37.8 Mb | 3.3 | Rice leaf folder | (Selvaraju et al. 2007) |
| **10** | 21.5 Mb | 22.9 Mb | 21.3 Mb | 22.7 Mb | 1.2 | Brown planthopper | (Sun et al. 2005) |
| **12** | 21.4 Mb | 23.3 Mb | 22.1 Mb | 23.2 Mb | 0.8 | Brown planthopper | (Jena et al. 2006) |

*^a^ this study.*

**Table S4:** List of differentially expressed Laccase-encoding genes that are situated in Chromosome 1 QTL interval.

| **Gene ID (MSU)** | **RAP_ID** | **SM_log_2_FC** | **SM92_log_2_FC** | **Annotation** |
| --- | --- | --- | --- | --- |
| LOC_Os01g61160 | Os01g0827300 | - | 7.15 | laccase precursor protein, putative, expressed |
| LOC_Os01g62480 | Os01g0842400 | - | 4.77 | laccase precursor protein, putative, expressed |
| LOC_Os01g62490 | Os01g0842500 | - | 9.58 | laccase precursor protein, putative, expressed |
| LOC_Os01g62600 | Os01g0843800 | - | 7.74 | laccase precursor protein, putative, expressed |
| LOC_Os01g63190 | Os01g0850700 | 1.31 | 1.38 | laccase precursor protein, putative, expressed |
